# Toward quantifying the adaptive role of bacterial pangenomes during environmental perturbations

**DOI:** 10.1101/2021.03.15.435471

**Authors:** Roth E. Conrad, Tomeu Viver, Juan F. Gago, Janet K. Hatt, Fanus Venter, Ramon Rosselló-Móra, Konstantinos T. Konstantinidis

**Affiliations:** Ocean Science & Engineering, School of Biological Sciences, Georgia Institute of Technology, Atlanta, Georgia, USA; Department of Biochemistry, Genetics and Microbiology, and Forestry and Agricultural Biotechnology Institute (FABI), University of Pretoria, Pretoria, South Africa; Marine Microbiology Group, Department of Animal and Microbial Biodiversity, Mediterranean Institutes for Advanced Studies (IMEDEA, CSIC-UIB), Esporles, Spain; School of Civil & Environmental Engineering, Georgia Institute of Technology, Atlanta Georgia, USA

## Abstract

Metagenomic surveys have revealed that natural microbial communities are predominantly composed of sequence-discrete, species-like populations but the genetic and/or ecological mechanisms that maintain such populations remain speculative, limiting our understanding of population speciation and adaptation to environmental perturbations. To address this knowledge gap, we sequenced 112 *Salinibacter ruber* isolates and 12 companion metagenomes recovered from four adjacent saltern ponds in Mallorca, Spain that were experimentally manipulated to dramatically alter salinity and light intensity, the two major drivers of these ecosystems. Our analyses showed that the pangenome of the local *Sal. ruber* population is open and similar in size (∼15,000 genes) to that of randomly sampled *Escherichia coli* genomes. While most of the accessory (non-core) genes showed low *in situ* coverage based on the metagenomes compared to the core genes, indicating that they were functionally unimportant and/or ephemeral, 3.49% of them became abundant when salinity (but not light intensity) conditions changed and encoded for functions related to osmoregulation. Nonetheless, the ecological advantage of these genes, while significant, was apparently not strong enough to purge diversity within the population. Collectively, our results revealed a possible mechanism for how this immense gene diversity is maintained, which has implications for the prokaryotic species concept.

**Significance Statement:** The pangenomes of bacterial species, i.e., the number of non-redundant genes carried by members of the species, can be enormous based on the genome sequencing of isolates from various sites around the globe and different years. However, to what extent this pattern of gene diversity applies to natural bacterial populations, i.e., strains co-occurring in the same site, and the value of this diversity for population adaptation during environmental transition remains unclear. This study showed that while the pangenome of a natural population can be similarly large, only a small fraction of the pangenome appears to be functionally important when conditions change. Taken together, these results provided quantitative insights into the extent and functional significance of the accessory pangenome of a natural, species-like population.

## Main

Our understanding of the intraspecific diversity of prokaryotes is based largely on the comparative analyses of collections of isolates. Since these isolates typically originate from a variety of samples, habitats, and times, they often show varying fitness backgrounds and genomic adaptations specific to the local conditions at the time and place of isolation. Accordingly, the number of nonredundant genes (i.e., the pangenome) within many of the species formed by such isolates appears to increase continuously with the addition of each new isolate (i.e., the pangenome is open), and thus is quite large e.g., >30,000 or more genes than the human genome. Pangenomes are comprised of core and accessory genes (1-4). Core genes are shared by all or almost all (>90% of the total) genomes of a species and account for the general ecological and phenotypic properties of the species. Accessory genes, also referred to as auxiliary, dispensable, variable, or flexible genes, are present in only one or a few genomes of a species and can be further divided into strain-specific (isolate-specific), rare, or common genes based on the fraction of genomes found to contain the gene. While this phenomenon is well documented, it is still unclear whether results from the comparison of isolates acquired from different habitats and samples translate well to the diversity within natural populations; that is, a population of conspecific strains co-existing in the same environment or sample.

The emerging picture from culture-independent metagenomic surveys of microbial communities is that bacteria and archaea predominantly form species-like, sequence-discrete populations with intraspecific genome sequence relatedness typically ranging from ∼95% to ∼100% genome-aggregate nucleotide identity (ANI) depending on the population considered i.e., populations having a more recent bottleneck or genome sweep event show lower levels of intraspecific diversity. In contrast, ANI values between distinct (interspecific) populations are typically lower than 90% (5-7). Sequence-discrete populations have been commonly found in many different habitats including marine, freshwater, soils, sediment, human gut, and biofilms, and are typically persistent over time and space (8-11) indicating that they are not ephemeral but long-lived entities. Indiscrete populations are rarely encountered in these previous studies, and when found are almost always attributable to the mixing of distinct habitats such as the mixing of water from different depths in the ocean water column during upwelling events (8, 11, 12). Consistent with the metagenomics perspective, a recent analysis of all available isolate genomes of named species (n ∼ 90 000) revealed a similar bimodal distribution in ANI values; that is, a small number of genome pairs show 85-95% ANI relative to pairs showing either >95% or <85% ANI (i.e., an “ANI gap” or “discontinuity”) (13). These data reveal that a similar genetic (sequence) discontinuity is characteristic of both naturally occurring populations as well as traditional “species” classified based on genomes obtained from pure cultures. It remains to be tested, however, if functional (gene content) diversity patterns are also similar between naturally occurring populations and pure culture collections. Furthermore, it is equally important to elucidate the gene content dynamics of local populations to better understand the underlying evolutionary processes that shape species-like, sequence-discrete populations and maintain coherent species-like genomic structure on a global scale [reviewed in (5, 14)].

More specifically, quantifying the extent of gene content variation (i.e., the accessory pangenome) within natural microbial populations is important to better understand the metabolic and ecological plasticity of a population and how accessory genes facilitate adaptation to environmental perturbations. One prevailing hypothesis is that non-core gene diversity is largely neutral or ephemeral resulting from random genetic drift and a lack of competition among members of the population that is strong enough to lead to complete dominance of the member(s) carrying the genes in question (15). A competing hypothesis is that co-occurring subpopulations may accumulate substantial and ecologically important (non-neutral) gene content differences that enable, for instance, differentiated affinity for the same substrate, and thus are on their way to parapatric or sympatric speciation (14, 16, 17). The experimental data to test these hypotheses rigorously and quantitatively for a natural population are currently lacking.

Metagenome-assembled genomes (MAGs) obtained from environmental samples using population genome binning techniques have been used to study sequence-discrete populations. However, verifying the purity, completeness, and accuracy of these MAGs is challenging (18-22). Furthermore, even with high quality MAGs, the extent to which the recovered gene content represents the pangenome of a population remains speculative. MAGs cannot fully capture the total standing gene content variation of a natural population due to i) limitations in short-read assembly of hypervariable or genomic repeat regions, ii) low coverage of rare or accessory genes iii) high coverage of conserved regions shared across multiple populations, and iv) challenges in accurately grouping assembled contigs into MAGs during population genome binning (11, 23, 24). Although a few longitudinal shotgun metagenomic studies have attempted to quantify the genetic variation within natural microbial populations, their primary focus has been on single nucleotide polymorphisms (SNPs) (e.g., allelic variation) rather than gene content variation (6, 25-27). A few studies have also shown that gene content can fluctuate within a population as an effect of the dominance of different strains (6, 11, 23) by querying isolate genomes or MAGs against time or spatial series metagenomes, but to our knowledge no study has quantified gene content diversity within natural populations or the existence of distinct (co-occurring) subpopulations based on shared gene content.

Solar salterns are semi-artificial environments used for harvesting salt for human consumption, and they harbor reduced microbial diversity driven primarily by environmental stressors, most notably sunlight intensity and high salt concentrations (28). *Salinibacter ruber* represents the major component of the bacterial fraction of salterns and is commonly isolated from hypersaline habitats globally (29). Hence, we used solar salterns in Mallorca, Spain as our experimental system, and focused on *Sal. ruber* to quantify the intraspecific gene content variation of its naturally occurring population on a local scale by combining metagenomic sampling with extensive isolate culturing efforts on the same samples. Samples were collected during a multi-stressor mesocosm experiment wherein salterns were stressed by different UV light and salinity exposure regimes. Specifically, one control and three experimental ponds were filled with the same pre-concentrated inlet brines (Fig. S1). Apart from the inlet brines coming from the same source, the experimental ponds were isolated (no brine flow between ponds) and challenged by: i) sunlight intensity alterations using a shading mesh to cover a previously unshaded pond and ii) uncover a previously shaded pond, resulting in ∼37-fold reduction or increase in sun irradiation, respectively (28), and, iii) abrupt changes in salt concentration from ∼34% (salt saturation or precipitation level) to ∼12% by dilution with freshwater over a period of four hours ((30), Fig. S1). The *Sal. ruber* population in the four ponds was followed for one month after these treatments, when salt saturation conditions had re-established in the dilution pond (iii) above due to natural evaporation, by recovering 207 *Sal. ruber* isolates and 12 whole-community shotgun metagenomes from three time points (one day, one week and one month post-treatment). Herein, we report the observed gene content diversity and the relative *in situ* gene abundance of the local *Sal. ruber* population during ambient (control) and experimentally altered environmental conditions.

## Results

### Sampling the local *Sal. ruber* population

To characterize the intraspecific diversity of the local *Sal. ruber* population *in situ*, we isolated 207 strains during a one-month time period from four adjacent saltern ponds at ‘ Es Trenc’ in Mallorca, Spain (Fig. S1). Based on MALDI-TOF MS and RAPD signatures (Fig. S2), we selected 123 non-clonal isolates (i.e., strains with different RAPD profiles) for genome sequencing. After genome assembly, we selected 112 draft genomes that were determined to be free of contamination and of good quality compared to previously completed *Sal. ruber* genomes for subsequent analyses (Sup. Excel File 1). Our genomes had an average of 267.53 (stdev=60.00) contigs per assembly, an average N50 value of 25,368.64 bps (stdev= 7,357.42) and an average sequencing depth of 14.51X (stdev=5.15). The mean genome sequence length was 3,828,264 bps (stdev=140,341.62) with an average of 65.83% G+C content (stdev=0.25%) and an average of 3,369 unfiltered open reading frames per genome. All the *Sal. ruber* draft genomes had one 16S rRNA gene copy of 1,535 bps in length.

### Genomic diversity of the local *Sal. ruber* population based on 112 isolates

Somewhat surprisingly, ANI vs. shared genome fraction analysis among all 112 genomes revealed a second closely related yet distinct population (n=10) around 95% ANI to the primary *Sal. ruber* population (n=102) (Fig. 1A). While this secondary cluster appears to be a divergent *Sal. ruber*-like clade based on our ANI analysis and maximum likelihood trees from 16S RNA and single copy protein-coding genes (SCGs) (fig 1A and S4), it was initially identified as *Sal. ruber* using MALDI-TOF MS and RAPD analyses due to the highly similar spectra that made the strains of the new lineage to appear scattered among the true *Sal. ruber* (Fig S2). Genomes of each population cluster share greater than 97.5% ANI among themselves with roughly a 3% ANI discontinuity and 5% difference in the shared genome fraction between them (Fig. 1A). We repeated this analysis using unassembled reads mapped to the assembled genomes in order to sidestep any potential biases resulting from assembly vs. assembly comparison of draft (incomplete) genomes containing hundreds of contigs and found essentially the same results (Fig. S3). We focused the remaining analyses on the primary *Sal. ruber* population due to its larger number of isolate genomes and the fact that the ten *Sal. ruber* genomes from the NCBI database fell within this primary population (Fig. 1A). Since the genomes from NCBI were isolated from various sites across the globe, they are representative of the broader species level diversity compared to the locally sampled population of isolates in our collection. The ANI values among the members of the primary population averaged approximately 98.0% and the shared genome fraction averaged approximately 80.0%, revealing that, while these genomes share high sequence identity, about 20% of the gene content differs in pairwise comparisons (Fig. 1A and S4). These findings revealed substantial intraspecific sequence (e.g., ANI) variation within the local population equivalent to that of the broader *Sal. ruber* species population based on the 10 available genomes from NCBI or other species with several sequenced representatives (13).

**Fig 1.**
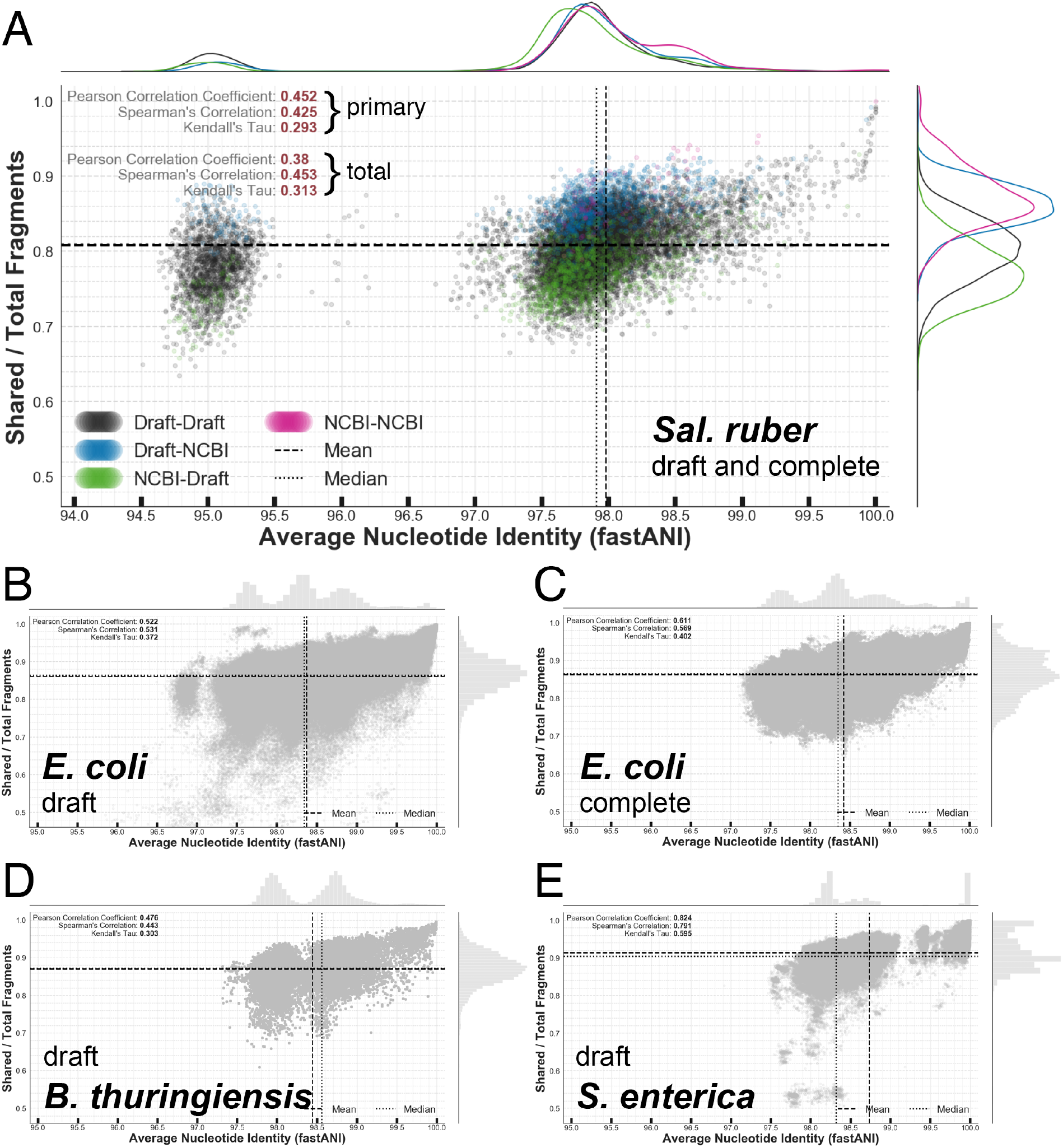
Genomic diversity within selected species assessed by ANI and shared genome fraction. The shared genome fraction (y-axis) is plotted against ANI (x-axis). Correlation coefficients between these two genome variables are shown on each plot along with lines for the mean (dashed) and median (dotted) values. The graphs to the top and right of each panel show (**A**) the kernel density estimates or (**B-E**) histograms for each variable. (**A**) Values representing the 112 x 112 comparisons of our *Sal. ruber* isolate draft genome collection (designated as Draft-Draft), 10 x 10 for *Sal. ruber* genomes from NBI (designated as NCBI-NCBI), and 112 x 10 or 10 x 112 for our draft genomes versus the complete NCBI genomes (designated as Draft-NCBI or NCBI-Draft, respectively). Correlation coefficients were calculated for the primary *Sal. ruber* population (primary) as well as for all the values (total). The means and medians were calculated for the primary *Sal. ruber* population only. Note that the draft-draft datapoints overlap with those datapoints representing complete genomes revealing no major biases by the draft nature of these genomes with respect to gene content or ANI values. (**B-E**) 100 x 100 genomic comparison results times 100 random sampling trials for (**B**) *E. coli* draft, (**C**) *E. coli* complete, (**D**) *B. thuringiensis* draft, and (**E**) *S. enterica* draft genome collections. Correlation coefficients, means, and medians were calculated for all values. Note the kernel density estimates (in A) and histograms (in B-E) on the perimeter of each panel (top and right) that clearly show the density distributions of the corresponding data points.

A maximum likelihood phylogeny of the full length 16S rRNA gene sequences carried by the isolate genomes or their concatenated set of 106 SCGs confirmed the ANI-based results. The two phylogenies showed that the minor *Sal. ruber*-like population formed a single diverging clade and that the ten NCBI genomes were dispersed throughout the primary clade (Fig. S4). In addition, the lack of defined subclades within the primary clade in terms of individual solar ponds (spatial) and time of sampling over the one-month sampling period (temporal) from which the genomes originated suggested that the local *Sal. ruber* population was homogeneous over space and time (Fig. S4). Likewise, the interspersed placement of the complete NCBI genomes within this primary clade indicated that genomic diversity at the local scale captured the diversity present at the global scale (Fig. S4). Collectively, these results indicated that our draft genome collection represents well the extant cultivatable genomic diversity found within the local *Sal. ruber* population and probably within the broader species population as well.

### Pangenome structure of the local *Sal. ruber* population

To assess genomic diversity at the gene level, we randomly selected 100 genomes from the primary *Sal. ruber* population (n=102) and analyzed the pangenome structure by calculating the empirical, non-redundant gene rarefaction curve while tracking new gene additions and gene class counts. We estimated the total pangenome (i.e., the number of non-redundant genes) of the local *Sal. ruber* population to be open (γ = 0.36) consisting of 12,666 genes in total (Fig. 2A upper panel). Each additional *Sal. ruber* genome added 98 new genes to the pangenome, on average, with a persistent mean addition of 48 new genes at n = 100 after 1,000 permutations of the order that genomes were added to the rarefaction curve (Fig. 2A, lower panel). The exponential decay model fit to these data estimated that the new gene ratio (number of new genes per genome added to the pangenome / number of genes in genome) reached a lower asymptotic value of Ω = 2.21% of genes per genome, although the empirical values were measured to extend below this with a mean value of Ω = 1.67% at n = 100. These data show that nearly 2% of the gene content in any *Sal. ruber* genome sampled by our collection consisted of unique genetic material and that because of this, the total gene content diversity remained under-sampled even after sampling 100 genomes collected from the same source water (Fig. 2A). Accordingly, we calculated that the pangenome of the local *Sal. ruber* genome collection was composed of 4,830 (∼38% of total) isolate-specific genes and 5,587 (∼44%) rare or common genes distributed among the members of the population, with the core genes making up (just) the remaining 18% (Fig. 2A).

**Fig 2.**
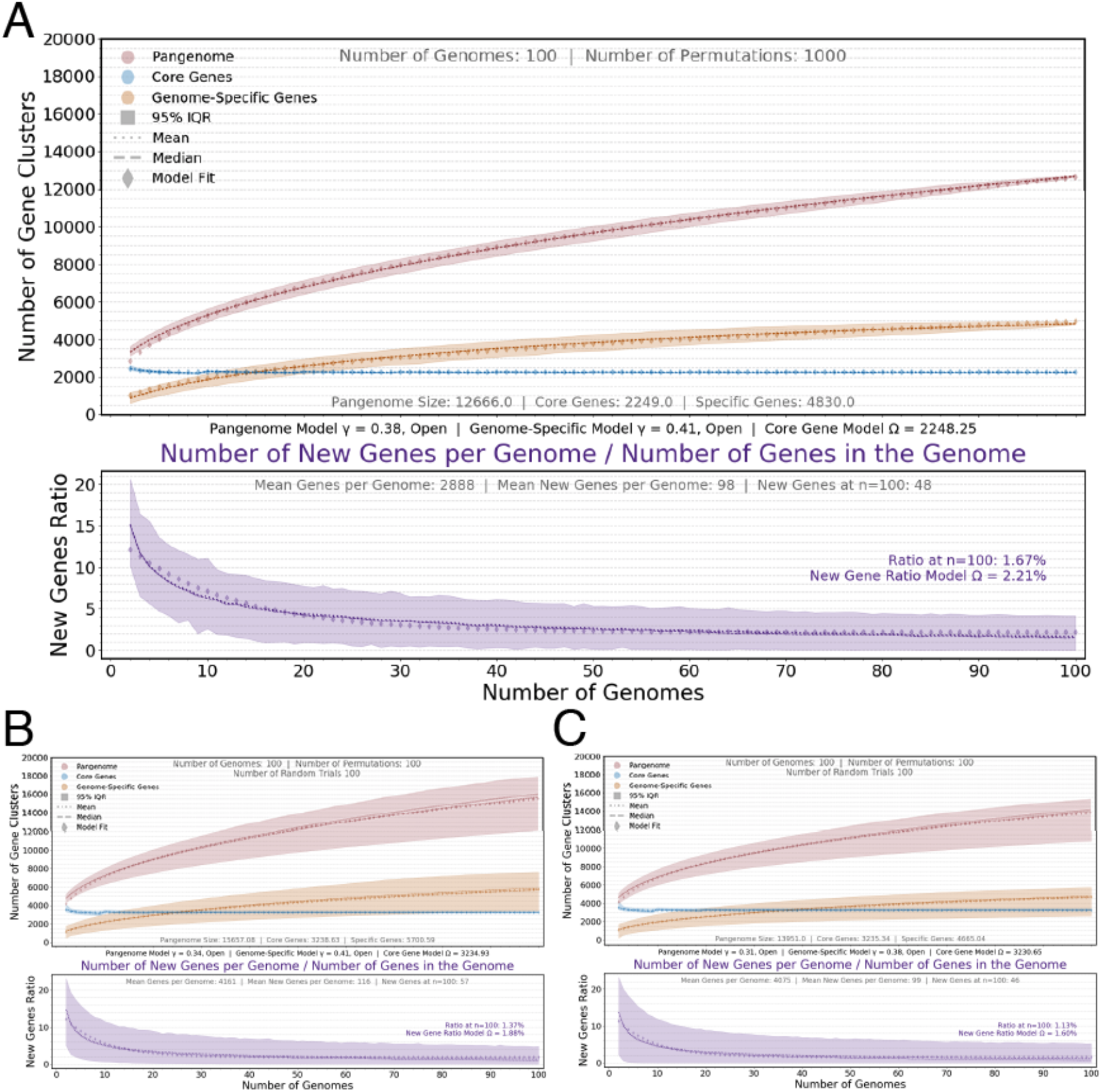
Pangenome comparison of *Sal. ruber* to selected bacterial species. (**A-C**) The top panel shows the mean, median, 95% empirical confidence interval of permuted values, and the model fit for each of three curves showing the total non-redundant genes in the pangenome (in red), total number of core genes (in blue), and total number of isolate-specific genes (in orange; y-axis) plotted against the number of genomes sampled (x-axis) for (**A**) 100 draft genomes from the primary *Sal. ruber* population, and results from all random trials for (**B**) *E. coli* draft genomes and (**C**) *E. coli* complete genomes. The lower panel shows the same calculations but for the new gene to genome ratio (in purple). The axes scales are conserved between panels.

While the total accessory genome remained unsaturated by sequencing, we estimated that there were 2,249 universally shared genes comprising the *Sal. ruber* core genome, or about 77.87% of a *Sal. ruber* genome (2,249 / 2,888) represented conserved, shared genetic material (Fig. 2A). These results were congruent with the ANI vs. shared genome fraction analysis as well (Fig. 1A). The persistence of the core genome was also validated by the extremely narrow empirical confidence intervals and by the exponential decay model reaching a similar lower asymptote limit at Ω = 2,248.25 after only 30 genome additions (Fig. 2A). Interestingly, genome clustering based on the presence and absence of accessory genes revealed a similar overall clade structure to the core gene phylogeny, although several genomes clustered differently between the two trees (Fig S5). Both trees indicated the existence of (coexisting) subpopulations within the *Sal. ruber* population based on the four or more distinct subclades recovered in the trees (Fig. S5). Consistent with this view, analysis using evolutionary read placement of metagenomic reads to reference *Sal. ruber rpoB* gene sequences indicated the presence of subpopulations (or genotypes) that fluctuated in abundance relative to each other across the different sampling times (Fig. S6).

### Pangenome structure compared to other model species populations

Using the same pipeline, with additional random trials to calculate empirical confidence intervals, we estimated the pangenome of several, phylogenetically and physiological diverse, model bacterial species including *Escherichia coli, Bacillus thuringiensis, Salmonella enterica, Mycobacterium tuberculosis*, and *Pseudomonas aeruginosa*. For these comparisons, we chose genomes to emulate the ANI distribution observed within the primary *Sal. ruber* population (∼98.0% ANI, on average) in order to avoid the known effect of higher gene content conservation among genomes with higher ANI (genetic) relatedness as noted previously (31). Our selection process generated intraspecific ANI distributions similar to the primary *Sal. ruber* population centered around ∼98.5% ANI (Fig. 1). We found the pangenome of *E. coli* to be open (γ = 0.34 for draft genomes and γ = 0.31 for complete genomes, Fig. 2 B, C) and similar in size to the primary *Sal. ruber* population pangenome (Figs. 2 & S7). Annotation results for the total pangenome were similar overall between *Sal. ruber* and *E. coli*, but, as expected for a well-studied model species, *E. coli* had fewer genes annotated as hypothetical or uncharacterized (Fig S7). Compared to *E. coli, Sal. ruber* had a comparable number of isolate-specific genes despite the smaller size of the *Sal. ruber* genome (2,888 vs. 4,118 genes per genome, on average). This was evident in a larger new gene ratio estimate for the *Sal. ruber* genomes, and in the number of isolate-specific (or rare) genes compared to the core genes (Fig. S7 C, E, F, H), especially after normalizing by the pangenome or genome size for each species (Fig. S7 D, I, J, L, M, N, P). Similar results to those reported for *E. coli* above were obtained for the other model bacteria species (not discussed further to avoid redundancy; results are available in Fig. S7).

Another interesting pattern revealed by our pangenome analysis was a consistent ratio of core genome size to genome size (i.e., what fraction of the total genes in the genome the core genes make up) at about 0.8-0.9, observed across species despite the variation of core gene to pangenome size ratios (Fig S7 P vs. L). This result was consistent with earlier observations based on a much smaller number of genomes per species (n ∼ 10) (16). In addition, there was a general lack of genes with intermediate prevalence in the pangenome of a species between the core and rare or isolate-specific gene classes (Figs. 3B & S7 E-P). This means that genes are either predominantly present in all genomes or in only a few genomes of the population in accordance with the idea that new beneficial gene sweeps are comparatively less common than the creation and subsequent loss of new genetic material. In any case, the *Sal. ruber* genomes seem to dedicate a larger portion of their genome to gene content variation compared to the other species analyzed here.

**Fig 3.**
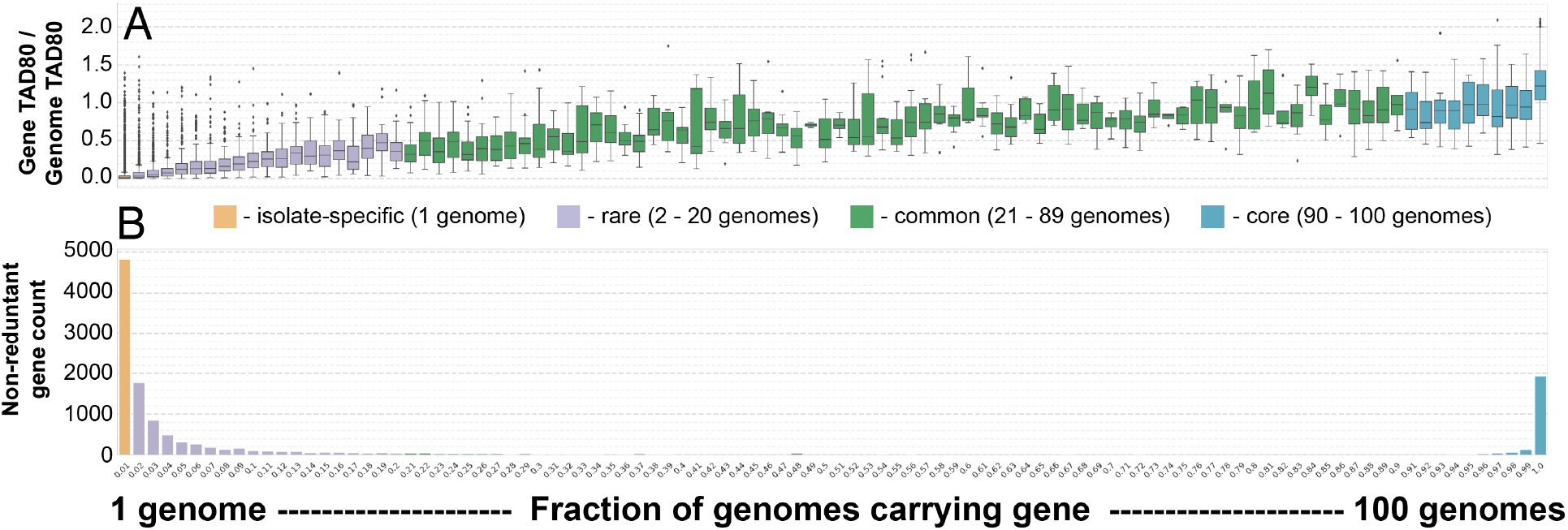
Relative *in situ* abundance of different classes of *Sal. ruber* genes based on their prevalence in isolate genomes. (**A**) Box plots show the distribution of the average TAD80 (i.e., *in situ* abundance) for each non-redundant gene cluster normalized by the average whole-genome TAD80 that was computed for and averaged across three control pond (salt saturation conditions) metagenomic datasets (y-axis). A normalized value of 1.0 indicates that the gene has an equivalent *in situ* coverage (abundance) to a single copy core gene. Genes are ordered on the x-axis based on their prevalence among 100 *Sal. ruber* genomes expressed as n / N where n ranges from 1 to N in increments of 1, and N = 100, e.g., a value of 0.01 (1/100) on the far left side indicates that only 1 genome has this gene cluster (isolate-specific gene class; orange), and values of 0.90 - 1.00 indicate that 90 to 100 genomes have the gene cluster (core gene class; blue) etc. (**B**) Counts of the number of non-redundant genes of the pangenome (y-axis) assigned to each gene prevalence class (x-axis); classes are ordered as in panel A.

### Ecological/Functional importance of rare and isolate-specific *Sal. ruber* genes

To test if any isolate-specific or rare genes could provide an ecological benefit or were instead functionally neutral and/or ephemeral, we assessed their relative *in situ* abundance in the companion metagenomes representing changes in environmental conditions. To estimate the sequence depth of each gene (coverage), we computed a truncated average depth using the middle 80% of the sequence base positions (TAD80) to remove outlier effects from short conserved domains or motifs and the edge effect of read mapping to the ends of contigs (i.e. the top and bottom 10% of per base sequence depth values are removed from the distribution prior to taking the average), as suggested recently (32). We then normalized the TAD80 for each gene cluster (Fig. S9) by the average whole-genome TAD80 (Fig. 3A) providing a view of the relative abundance of genes in relation to the relative genome abundance. Thus, a normalized value of 1.0 indicated that a gene has an equivalent *in situ* sequencing depth (relative abundance) to a single copy core gene; a value above 1.0 indicated that the gene abundance is greater than the genome average. The resulting data revealed a strong decreasing trend in the distribution of gene abundances from the core gene class to the isolate-specific gene class (Fig. 3A). Notably, we identified that 0.66% of the isolate-specific genes (i.e., present in only one genome in our collection) and 2.83% of the rare genes (i.e., present in <20% of the genome in our collection) became abundant *in situ* during the intermediate salinity conditions (diluted pond one-week sample; 23.6% salt concentration), which followed the dilution from high (∼34% salts) to low (∼12% salts) salinity (Fig. 3). Notably, the *Sal. ruber* population abundance in these samples did not vary more than three-fold (Fig. S9); thus, the differential gene abundance reported above cannot be attributed to possible artifacts related to low sequence coverage of the population.

Functional annotation of the abundant fraction of isolate-specific and rare genes from the intermediate-salinity metagenome revealed that several of these genes could be involved in response to osmolarity changes, gene regulation, and transport of metabolites in/out of the cell. (Fig. 4H). While this trend was also observed for a few genes in the other gene classes (Fig. 4 B, E), several of the isolate-specific genes that peaked in abundance during intermediate salinity shared high sequence similarity to genes found on the pSR84 plasmid from *Sal. ruber* strain M8, isolated more than a decade ago and shown to be more tolerant of lower salinity conditions than the type strain of *Sal. ruber* (Strain DMS 13855 or M31) (33). These results contrasted with an over-dominance of hypothetical and mobile functions among the isolate-specific genes that did not change in abundance in the intermediate-salinity metagenomes (Figs. 4 & S10), revealing a strong bias toward functions that are presumably related to the salinity perturbation.

**Fig 4.**
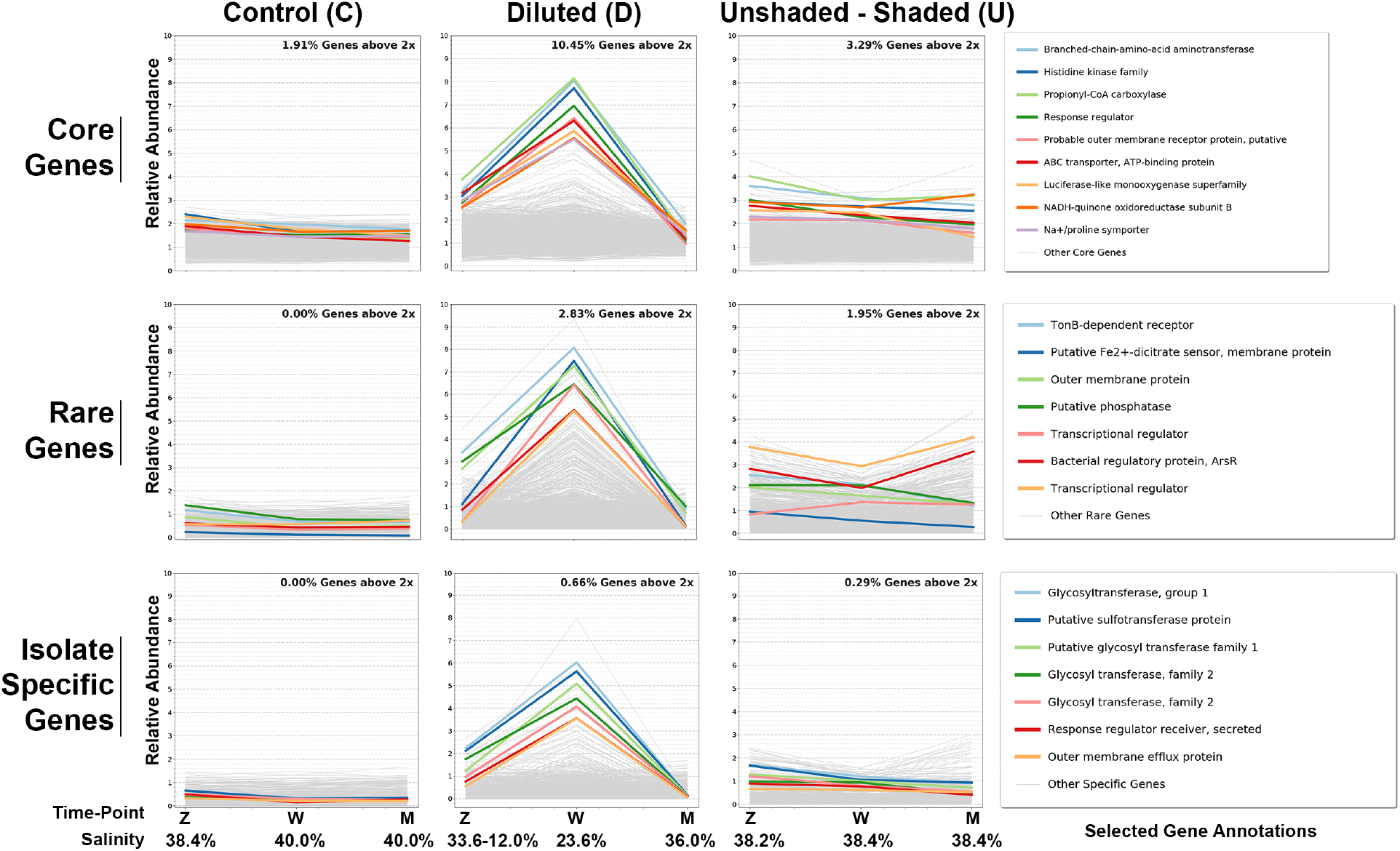
*Sal. ruber* gene abundance dynamics over the one month period of the experiment. Each line represents a non-redundant gene of the *Sal. ruber* pangenome for which the average TAD80 of the gene normalized by the whole-genome TAD80 of the same sample (y-axes) is plotted against the three metagenomic sampling time points (x-axes) for each of the three separate experimental ponds outlined in Fig. S1. Therefore, the lines represent the relative gene abundance in relation to the relative genome abundance. Results are organized by core (**A-C**), rare (**D-F**), or isolate-specific (**G-I**) gene class. Results for all genes are plotted as gray lines with a handful of genes from each class, selected based on their (higher) peak in **B, E**, and **H** panels representing the pond with changing salinity conditions (see x-axis values), shown in color. The corresponding functional annotations of these genes are also shown (figure legend).

In addition, 10.45% of the core genes also showed increased relative *in situ* abundances at the intermediate salinity metagenome and the (predicted) functions encoded by these core genes (Fig. 4B) were similarly involved in osmoregulation and transport as the rare (Fig. 4E) and strain-specific (Fig. 4H) genes mentioned above. Hence, the enrichment of (or selection for) specific functions during salinity transition was evident in different parts of the population’ s pangenome, and we focused our analysis on strain-specific and rare genes because these made up a larger fraction of the pangenome (Fig. 3). Given that core genes are typically single-copy genes in the genome, the increased abundances noted (Fig. 3B) could be due to recent horizontal transfer of these genes to/from other co-occurring populations or duplication of the genes within the genome (e.g. they are carried by multi-copy plasmids). Future work will elucidate the relevance of each of these scenarios.

Finally, we assembled our metagenomic samples to evaluate the diversity of the genomic background (origin) of the *Sal. ruber* isolate-specific genes that became abundant under the low-salinity condition. We found a range of contigs within each metagenome that showed high identity matches, in the 70 – 100% sequence identity range, to these genes (Fig. S11 & S12), but these contigs also contain other genes that do not match the *Sal. ruber* genomes. These results revealed that additional community members encode the genes, especially in the dilution pond one-week time point sample (salinity 23.6% NaCl), indicating that the corresponding functions may indeed be selected by intermediate salinities. In general, these contigs also show gene synteny among themselves, indicating a common origin for the corresponding genes and operons (Fig. S13).

## Discussion

The pangenome of a local *Sal. ruber* population, sampled over a one-month period from four adjacent saltern ponds filled with the same source water, is open and similar in size to the pangenomes of *E. coli* and other species whose genomes were recovered from around the globe over the course of many years (Fig. 2). These results were somewhat unexpected given the range of samples providing the *E. coli* (and other) genomes relative to the few samples that provided the *Sal. ruber* isolates. In fact, we found that the pangenome to genome size ratio of the local *Sal. ruber* population is the largest of all species considered (Fig. S7B). While pangenome sizes are known to vary between species (15, 17, 34), it is intriguing to find such extensive gene content variation within a single population and local source. These results also corroborate observations from a previous study of another large *Sal. ruber* isolate collection retrieved from a single, one drop (0.1 ml) sample that showed *Sal. ruber* to be phylogenetically homogenous at the ribosomal level yet diverse based on restriction patterns and metabolomics analysis (35). Furthermore, the great majority of non-redundant *Sal. ruber* genes were either isolate-specific, very rare or core genes (Figs. 3B & S7E-P) and few genes were found at intermediate prevalence in the primary *Sal. ruber* population. These results imply that the majority of non-core genes are indeed neutral and/or ephemeral as previously hypothesized and do not contribute to the major functions carried by the population. Horizontal gene transfer (HGT) and gene deletion presumably underlie these patterns. Consistent with this assumption, our comparison of the *Sal. ruber* SCGs phylogenetic tree to that of the dendrogram based on the presence or absences of gene content indicated that HGT (and gene deletion) may be frequent (Fig. S5).

Interestingly, while the great majority of isolate-specific and rare genes remained rare as conditions changed, at least a few of them (about 3.49% of the total) were found to considerably increase in abundance during the low-salinity (dilution) perturbation. Several of the latter genes were three times more abundant relative to the genome average at intermediate salinity. Notably, the predicted functions encoded by these genes were associated with environmental sensing, metabolite transport in/out of the cell, glycosyltransferases (which may be related to osmoregulation) (36), and gene regulation (Figs. 4, S11, S12, S13). Very few hypothetical or uncharacterized genes were identified among the genes showing increased abundance at intermediate salinity despite the high frequency of the former genes in the total pangenome (Fig. S10). Collectively, these results further supported our hypothesis that the identified genes involved in regulation and transport are presumably important for cell osmoregulation under low- and intermediate-salinity conditions. While these findings await experimental testing, they do indicate that the isolate-specific and rare genes identified may facilitate the *Sal. ruber* population in adapting to changes in salinity concentrations, and thus are likely not neutral or ephemeral. Unfortunately, the exact function or substrate specificity for the identified genes remains unknown as bioinformatics analysis provides only general functional prediction. Hence, we are not yet able to make specific inferences about how exactly these genes facilitate adaptation to intermediate salinity conditions. Note also that assessing the relative abundance of these genes in the time zero sample of lowest salinity (∼12% salts) would not have been meaningful in this respect because the corresponding cells that carry these genes did not have enough time to adjust to the low salinity conditions and begin to grow (37). Further, the great majority of the identified genes apparently do not undergo adaptive evolution since their pN/pS ratio based on metagenomic reads mapped on the genes were low, between 0.1-0.5, although slightly higher than that of the core genes (Fig. S14). These ratios indicated strong purifying selection for the identified osmoregulation-related genes and that they are already well fit for the function they carry out.

It is possible that additional isolate-specific or rare genes become important during fluctuations of other environmental variables. However, analysis of metagenomic data from the experimental manipulation of light intensity, the other major environmental factor for the saltern ecosystems (28), did not reveal any isolate-specific genes to change in abundance as we observed in the intermediate salinity samples. Thus, any additional isolate-specific or rare genes of ecological importance would have to be specific to environmental parameters not measured by our work. Such parameters could include seasonal fluctuations (e.g., we sampled for one-month in August) or biotic factors such as phage predation (38). In any case, genes of such ecological importance are not expected to make up a large fraction of the pangenome based on the results reported here (e.g., most isolate-specific genes were not found to be abundant relative to the average genome relative abundance in any metagenomic datasets) for salinity and light intensity transitions. While some of these results and interpretations echoed those in previous studies (16, 39-41), they do provide a new and more quantitative perspective into the role of biodiversity within sequence-discrete populations (and species) during environmental transition.

An emerging question based on these findings is why the intrapopulation diversity was not purged (removed) when salinity conditions changed. That is, the genomes (cells) that encode the abovementioned genes should have outcompeted the remaining genomes of the population resulting in a more clonal population and/or (sub-population) speciation. However, phylogenetic analysis of the isolates (Fig. S2) and metagenomic read placement (Fig. S6) suggests that intrapopulation diversity was maintained and, in fact, the dominant subpopulations in terms of gene content bounced back in abundance when salt-saturation conditions were re-established. Thus, we hypothesize that the ecological advantage of these genes is significant, but not strong enough to purge the intrapopulation diversity (or sweep through the population) or a much longer duration of intermediate salinities than represented by our experimental design or the typical salt cycles observed in the Mallorca salterns would have been required for a population sweep event to take place. Consistent with this working hypothesis, genomes that do not encode these genes were apparently able to survive at lower growth rates until favorable (salt-saturation) conditions returned (Fig. S6). While this hypothesis remains to be experimentally tested, it does provide a plausible mechanism that would maintain sequence-discrete populations despite such immense intrapopulation gene content diversity, frequent HGT, and environmental transitions. Moreover, these results further corroborate the use of sequence-discrete populations as the important unit of microbial diversity for taxonomy as well as for future investigations for advancing taxonomy and diversity studies.

It is challenging to capture the complete gene content of natural populations based on short-read metagenome sequencing, e.g., the average bacterial gene is one Kbp in size which is much longer than the typical length of short Illumina reads (150-250bp) that predominate recent metagenomic studies. Furthermore, it is unlikely that all accessory genes can be recovered by MAGs [discussed above and in (20, 21, 42)]. While isolate sequencing can circumvent these limitations, isolation may provide an uneven view of the natural population due to isolation biases. However, this is unlikely to have been the case for the primary *Sal. ruber* population studied here based on several independent lines of evidence. First, the ten *Sal. ruber* genomes from NCBI, which originated from various salterns around the world, are grouped together into a single clade with the 102 genomes of the primary population in our core-genome phylogeny. This is also the case for the evolutionary placement of metagenomic reads onto the genome-based phylogeny, i.e., the great majority of reads were assigned to terminal branches (tips) of the tree as opposed to ancestral nodes; the latter would have indicated that the reads originated from abundant strains not represented by our isolates. Second, our collection of sequenced isolates represents a much larger collection (n = 207) of isolates that were first identified as *Sal. ruber* by MALDI-TOF MS whole cell analysis and dereplicated by RAPD profile analyses to avoid sequencing the same clone. Importantly, in this larger collection, there were not any major or minor subclade(s) that were not represented by the subset of the 112 isolates we sequenced (Fig. S2A). Hence, our genome collection appears to be representative of the natural *Sal. ruber* population based on the phylogenetic placement of metagenomic reads, the MALDI-TOF spectra analysis, and the RAPD profiles of a much larger collection of (non-sequenced) isolates (Fig. S6 and S2). Finally, based on the ANI vs. the shared genome fraction analysis (Fig. 1), the draft *E. coli* genome collection had a wider distribution than the complete *E. coli* genome collection; however, the median values on both axes were nearly equal (Fig. 1 B & C). The wider distribution could be a technical artifact due to the nature of incomplete versus complete genome sequences, or likely a true signal of biological diversity arising from the greater number of draft genomes available. Regardless of the exact underlying reason, the similarity in values indicates that our measurements from draft genomes provide similar estimates to complete genomes and that our pipeline was robust. Importantly, we selected draft genomes from different species to be of a similar level of completeness and ANI relatedness to avoid any systematic effect of these parameters on our results and conclusions.

Despite the abovementioned advantages, the number of genomes we sequenced was still limiting compared to the total gene content diversity as evidenced by the incomplete capture of accessory genes by our pangenome analysis. Future work could include more replicate samples, longer time series analysis, and deeper metagenomic sequencing with long-read technology for more robust results and interpretations. It would be particularly interesting to measure the fitness advantage of isolates based on their specific complement of accessory genes to directly test the hypotheses presented above related to the ecological advantage of such genes. The work presented here should serve as a guide for the number of samples and isolates to obtain, amount of sequencing to apply (Fig. S9), and what bioinformatics analyses to perform for studying the value of the diversity within natural sequence-discrete populations.

## Materials and Methods

### Experimental site, sampling and processing

*Sal. ruber* isolates and whole-community samples for shotgun-metagenomics were collected concomitantly from four adjacent solar saltern ponds in Mallorca, Spain at ‘ Es Trenc’ (Fig. S1) at three time points over a one-month period post-treatment in August of 2012. The saltern ponds are part of a larger facility of crystallizers for salt harvest and are fed with the same inlet brines. After filling with inlet brines there was no fluid exchange between the ponds during the course of study, and there were negligible effects from rainfall in Majorca, Spain during the month of August, when the experiment was conducted. The conditions for each pond were as follows: 1) a control pond with ambient sunlight and salt-saturation conditions found at the ‘ Es Trenc’ salterns, 2) a shaded-unshaded pond that was covered with a mesh to reduce sunlight intensity by 37-fold for 3 months prior to the experiment and then uncovered (unshaded) at time zero, 3) an unshaded-shaded pond that was kept uncovered until time zero, then covered with a mesh (shaded), and 4) a diluted pond to which freshwater was added to reduce the salinity from ∼34% to ∼12% in less than one hour. The mesocosm experiment was designed to test the effects of light intensity and salinity levels on the indigenous microbial communities inhabiting the salterns. Ponds #2 and #4 reached salt saturation conditions after one month during which time the microbial community dynamics re-stabilized; thus, no further sampling was performed after one month. The isolates used in this study were collected from all four ponds at time zero (just before the stressors were applied) and one-month (at the end of the experiment). Isolates were also sequenced from samples taken at two days post dilution treatment and one-week post the unshaded-shaded treatment (Fig. S1). The companion metagenomes for this study were sequenced from all four ponds sampled at one day, one week and one month post treatment (Fig. S1). Metagenomes for the shaded-unshaded experiment were excluded in this study due to low sequence coverage of the *Sal. ruber* population in the corresponding samples. The experimental design and detailed procedures were previously described in (28), except for the inclusion of the diluted pond experiment which is described in (30) and outlined in Fig. S1. Samples were collected, processed for culturing, and resulting isolates were identified using Matrix-Assisted Laser Desorption Ionization–Time of Flight Mass Spectrometry (MALDI–TOF MS) as described by (28, 29, 43) (Fig. S2A). Multiple clonal isolates were dereplicated using random amplified polymorphic DNA (RAPD) fingerprinting (43) (Fig. S2B).

### DNA extraction and Illumina sequencing

*Sal. ruber* isolate cultivation and DNA extraction were performed as described in (44, 45). For metagenomic DNA extraction, 25ml of brine samples were centrifuged at 13,000 rpm as detailed in (45). DNA sequencing libraries were prepared using the Illumina Nextera XT DNA library prep kit according to manufacturer’ s instructions up to the isolation of cleaned double stranded libraries. Library concentrations were determined by fluorescent quantification using a Qubit HS DNA kit and Qubit 2.0 fluorometer (ThermoFisher Scientific) and samples were run on a High Sensitivity DNA chip using the Bioanalyzer 2100 instrument (Agilent) to determine library insert sizes. Libraries were sequenced for 500-cycles (2 x 250-bp paired-end) on an Illumina MiSeq instrument (Molecular Evolution Core facility, Georgia Institute of Technology) as recommended by the manufacturer. Additional sequencing of selected low-coverage libraries after the MiSeq sequencing was carried out on an Illumina NextSeq 500 instrument (located in the same facility) using a rapid run of 300 cycles (2 x 150-bp paired-end). Adapter trimming and demultiplexing of sequenced samples was carried out by the software available on each respective sequencing instrument.

### Illumina sequence quality control, assembly, and gene prediction

Raw Illumina reads in fastq format were evaluated with FastQC version 0.11.2 (46) in addition to quality analysis using custom Python scripts. Trimming and adapter clipping were performed using Trimmomatic version 0.39 (47) with settings ILLUMINACLIP NexteraPE-PE.fa:2:30:10:2:keepBothReads LEADING:3 TRAILING:3 MINLEN:36. Assembly was performed using SPAdes version 3.13.0 with the “--careful” flag and “-k 21,33,55,77,99,127”. Gene prediction was performed using Prodigal version 2.6.3 with default settings (48). The resulting summary tables can be found in Supplementary Excel File 1.

### Assessment of draft genome quality and phylogenetic analyses

The *Sal. ruber* isolate draft genomes were assembled from an average of 157.64 Mbps (stdev=66.15) of Illumina sequenced reads per isolate after adapter clipping and quality trimming (Supplemental Excel File 1). For each assembly, contigs shorter than 1,000 base pairs with supporting sequence coverage of less than 2X were removed from the assembly. The draft genomes in addition to *Sal. ruber* and *Sal. altiplanensis* genomes retrieved from NCBI were evaluated using the Microbial Genomes Atlas (MiGA) (49) to generate assembly metrics, quality scores, and all vs. all ANI scores (Supplemental Excel File 1). MiGA also identifies and extracts predicted 16S rRNA gene sequences and universal SCGs for each genome submitted. The sequences for the 16S rRNA, rpoB and the concatenated set of SCGs from each genome were aligned using Clustal Omega version 1.2.1 (50) with default settings. Maximum likelihood trees for the 16S and rpoB alignments were generated using RAxML version 8.0.19 (51) with parameters: *-m GTRGAMMA -f a -N autoMRE -p 4564821 -T 2 -x 1235*. An approximate maximum likelihood tree was generated for the concatenated SGCs using FastTree v2.1.10 (52) with default settings. The trees were drawn using either FigTree v1.4.3 (53) or iTOL version 4 (54).

### Pangenome analysis

A custom pipeline was developed for the pangenome analysis using a combination of Bash and Python programming. The pipeline starts with a directory containing genomes for a single species and proceeds in five parts. <u>Part 1</u> selects a seed genome at random and then continues random selection of genomes without replacement until the requested number of genomes meeting the criteria is reached. The pipeline keeps a genome if it matches the seed genome above a user defined ANI value (97.5% by default to match the ANI values observed among *Sal. ruber* isolate genomes; Fig. 1). <u>Part 2</u> predicts genes for each genome selected using Prodigal and removes genes shorter than 300 nucleotides in length. <u>Part 3</u> runs an all vs. all AN genome comparison using FastANI version 1.1 (13). <u>Part 4</u> clusters all genes from all genomes using CD-HIT-EST version 4.7 (55) with parameters: *-c 0*.*9 -n 8 -G 0 -g 1 -aS 0*.*7 -M 10000 -d 0 -T 10*. <u>Step 5</u> uses custom python scripts to parse the CD-HIT cluster file, run permutations to calculate the pangenome statistics, fit models to the empirical data, and build graphical plots of the results. The genomes used in our analysis included 878 complete and 11,167 draft *E. coli*, 433 draft *Bacillus thuringiensis*, 3,037 draft *Salmonella enterica*, 1,865 draft *Mycobacterium tuberculosis*, and 3,264 draft *Pseudomonas aeruginosa* genomes downloaded from NCBI on June 26, 2019. To generate empirical distributions, the custom pangenome pipeline was used to run 100 random bootstrap trials for each species (other than *Sal. ruber*) with genome replacement between trials. Only one trial was run for *Sal. ruber* since only about 100 draft genomes were available and each trial used 100 genomes.

In accordance with Tettelin et. al. 2005 & 2008 (1, 3), we fit a power law model to our data to estimate the upward trajectory of the pangenome growth curve using the γ parameter, and, we fit an exponential decay model to our data to estimate the Ω parameter which represents the lower boundary of the curves for the number of core genes or new gene additions. The possible values for the γ parameter reflect an open (0 < γ < 1) or closed (γ < 0) pangenome. For this analysis, we chose 100 genomes at random from the primary *Sal. ruber* population which excluded the two draft genomes labelled as SZ05 and SM11 corresponding to the 5^th^ and 11^th^ isolates obtained from the shaded pond at time zero and one month respectively. We then defined gene classes by a parameter p = n / N where n is the number of genomes carrying a gene and N is the total number of genomes (N = 100). Core genes are defined as those showing p ≥ 0.9, common when 0.2 ≤ p < 0.9, rare when 1 / N < p < 0.2, or isolate-specific when p = 1 / N. Accordingly, the accessory pangenome consists of all isolate-specific, rare, and common genes.

### Estimation of *in situ* gene abundance

Quality-trimmed metagenomic reads were searched against *Sal. ruber* draft genomes separately for each genome using the blastn option from Blast+ version 2.2.29 with default settings (56). Reads that found a match higher than the 95% sequence identity threshold and an (alignment length) / (query read length) greater than a threshold of 0.9 were used to calculate sequence depth (relative abundance). The resulting read depth data was truncated to the middle 80% (TAD80) of depth values (i.e., the upper and lower 10% of outliers were removed) using a custom Python script to provide TAD80 values for each genome, contig, and gene (Figs. S9, 3, 4; Supplemental Excel File 2). The 2^nd^ pond (shaded-unshaded) had relatively lower *Sal. ruber* abundance compared to the other ponds presumably due to the long-term shading of the pond, and was sequenced at a lower effort, which rendered the assessment of gene *in situ* abundance unreliable. Hence, this pond was used for isolation but not in the remaining bioinformatics analysis.

### Gene annotations

Representative genes for each CD-HIT-EST gene cluster were annotated against both UniProt databases (SwissProt and TrEMBL release-2018_05) using Blastp with default settings. Results were filtered for best match using a minimum threshold of 40% sequence identity and 50% alignment length coverage of the UniProt sequence for a match. Genes were also annotated using KofamKOALA version 2019-07-03 (57) with KEGG release 91.0 (58) and only annotations with an asterisk indicating they were above predefined thresholds for the corresponding HMM models were kept for analysis. Annotations can be found in supplemental Excel file 2 and 3.

## Code Availability

The custom code for these analyses are available on GitHub: https://github.com/rotheconrad/Salinibacter_ruber_01.

## Data availability

The data for this study has been deposited in the European Nucleotide Archive (ENA) at EMBL-EBI under accession number PRJEB27680 (https://www.ebi.ac.uk/ena/data/view/PRJEB27680).

## Acknowledgements

The authors would particularly like to thank especially the whole team at Salinas d’ Es Trenc and Gusto Mundial Balearides, S.L. (Flor de Sal d’Es Trenc) for allowing access to their facilities and for their support in performing the experiments. This work was partly funded by the US National Science Foundation, awards #1831582 and #1759831 (to KTK), and by the projects CLG2015_66686-C3-1-P, PGC2018-096956-B-C41 and RTC-2017-6405-1 of the Spanish Ministry of Science, Innovation and Universities (to RRM), which were also supported with European Regional Development Fund (FEDER) funds. RRM acknowledges the financial support of the sabbatical stay at Georgia Tech by the grant PRX18/00048 also from the Spanish Ministry of Science, Innovation and Universities. We thank Miguel Rodriguez-R and Carlos Ruiz for useful discussions on the methodology applied and ecologic implications of our results as well as the Partnership for an Advanced Computing Environment (PACE) at the Georgia Institute of Technology, which enabled the computational tasks associated with this study.

