## Supplemental Figures for "Toward quantifying the adaptive role of bacterial pangenomes during environmental perturbations"

**Running Title**

The role of the pangenome in adaptation

**Authors Full Names and Affiliations**

Roth E. Conrad[1,5], Tomeu Viver[3,5], Juan F. Gago[3], Janet K. Hatt[4], Fanus Venter[2], Ramon Rosselló-Móra[3*], Konstantinos T. Konstantinidis[4*]

[1]Ocean Science & Engineering, School of Biological Sciences, Georgia Institute of Technology, Atlanta, Georgia, USA

[2] Department of Biochemistry, Genetics and Microbiology, and Forestry and Agricultural Biotechnology Institute (FABI), University of Pretoria, Pretoria, South Africa

[3]Marine Microbiology Group, Department of Animal and Microbial Biodiversity, Mediterranean Institutes for Advanced Studies (IMEDEA, CSIC-UIB), Esporles, Spain

[4]School of Civil & Environmental Engineering, Georgia Institute of Technology, Atlanta Georgia, USA

[5]These authors contributed equally to this work.

**Corresponding Author Info**

### SI Figures

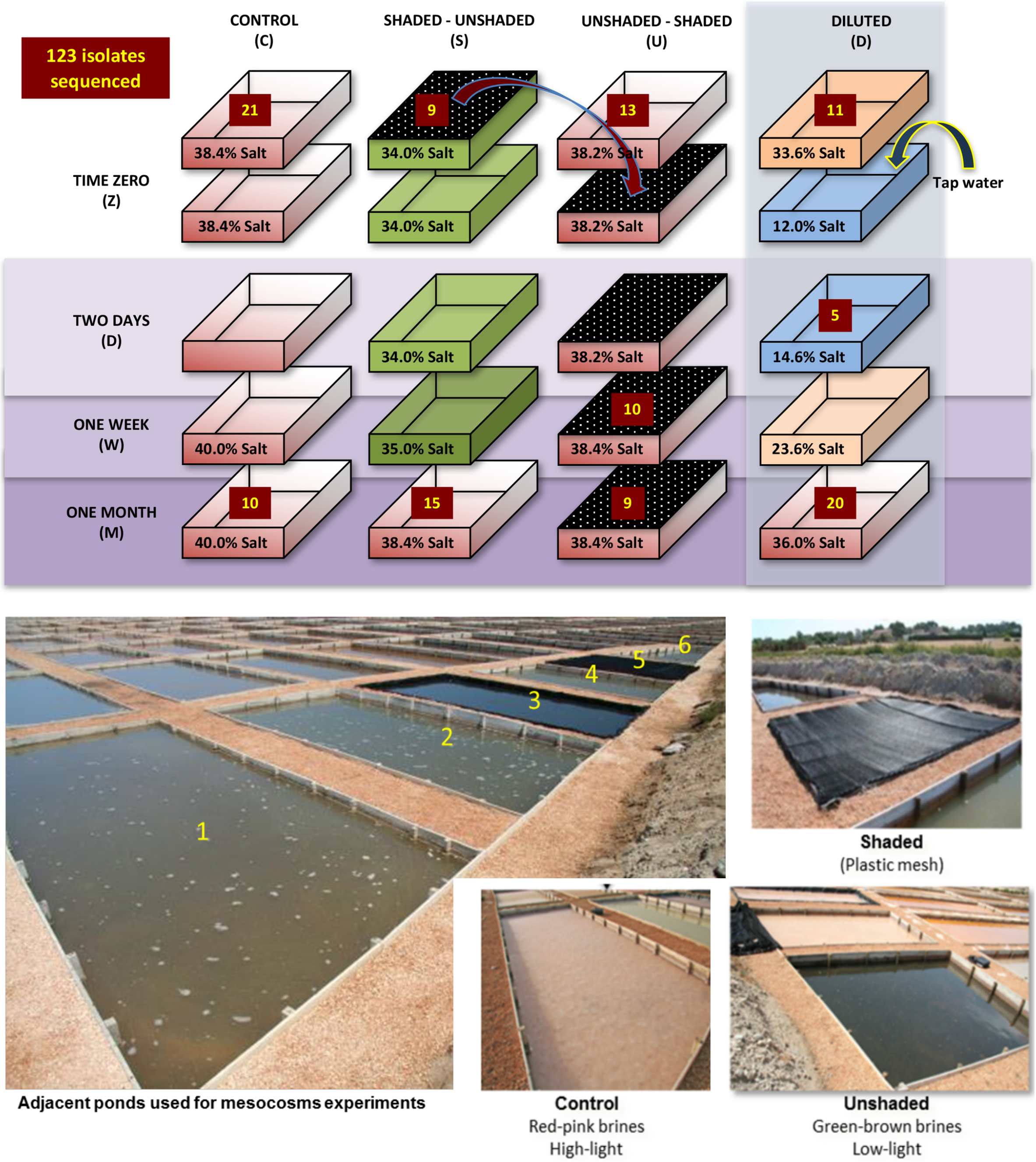

**Supplementary Figure 1: Experimental design.** Three saltern ponds at 'Es Trenc' in Mallorca, Spain were filled with the same pre-concentrated inlet brines in May of 2012. The brines in each pond were allowed to evaporate over the summer months with weekly refilling to reach NaCl saturation (>36% salts at 25˚C) as reported previously (1). The experiment was carried out in August 2012. Control and Unshaded-Shaded ponds were treated under standard operating procedures and environmental conditions of the site. The Shaded-Unshaded pond was covered with a mesh to reduce sunlight intensity by 37-fold in May and uncovered (unshaded) at time zero in August. Salinity for this pond was below saturation (34% Salts) at time zero and increased to saturation during the one-month experiment (1). The Unshaded-Shaded pond was covered with the same mesh to reduce light intensity at the start of the experiment. To avoid saturation and salt precipitation, the fourth saltern pond (Diluted) was pre-filled with the same pre-concentrated inlet brines two weeks prior to time zero and allowed to stabilize. Diluting a pond that has salt precipitate would require strong mechanical mixing of the brine to dissolve the precipitate. Hence, to avoid this complication, the salt concentration in the dilution pond was maintained, as close to salt saturation as possible without causing salt precipitation, by filling it with pre-concentrated brine two weeks before the onset of the experiment. On the day of the experiment, this pond was diluted by 2.8-fold using tap (freshwater) and then left to evaporate in the standard environmental conditions of the site. Salinity values given on the boxes indicate the salt concentration at the sampling times at which metagenomes were collected and culturing was performed. The numbers provided at each time point indicate the number of isolates from each sample that were sequenced for genome assembly. Companion metagenomes were sequenced for Control (C), Dilution (D), and Unshaded-shaded (U) ponds at times Zero (Z), one week (W), and one month (M). Metagenomes for the shaded-unshaded (S) pond were excluded due to low sequence coverage of the *Sal. ruber* population in these samples.

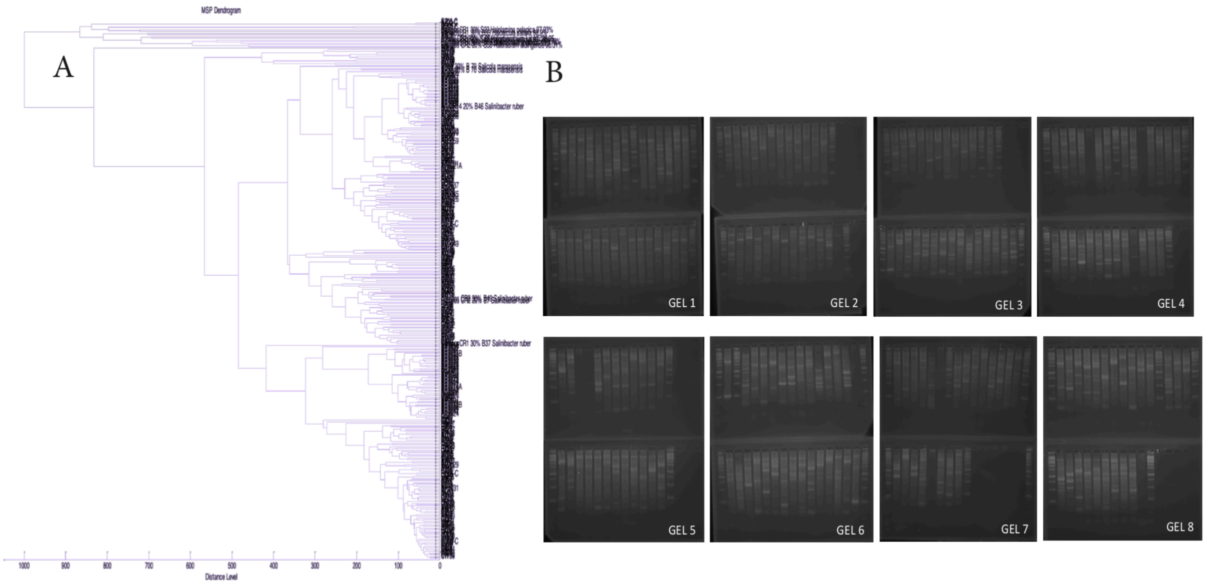


**Supplementary Figure 2: Identification of *Sal. ruber* isolates using MALDI-TOF MS and dereplication of clonal isolates using RAPD profiles.** Using the previously described cultivation approach in 25% saltwater medium (2), we isolated a total of 207 *Sal. ruber* isolates from the four saltern ponds. (**A**) To identify which of our cultured isolates were *Sal. ruber*, we generated whole cell MALDI-TOF MS profiles (3) of our isolates and compared the results to those of known *Sal. ruber* and other related species. The dendrogram shows results of a MALDI-TOF MS Spectra analysis of isolates collected for this study compared to an in-house profile database containing profiles from members of the *Salinibacteraceae* family and was generated using the BioTyper software 3.0 (Bruker Daltonics). (**B**) RAPD profiles (2, 3) were used to dereplicate clonal isolates to maximize sequencing effort. Identical RAPD profiles are interpreted to belong to clonal varieties. Each column in the gels shows a RAPD profile for a different isolate.

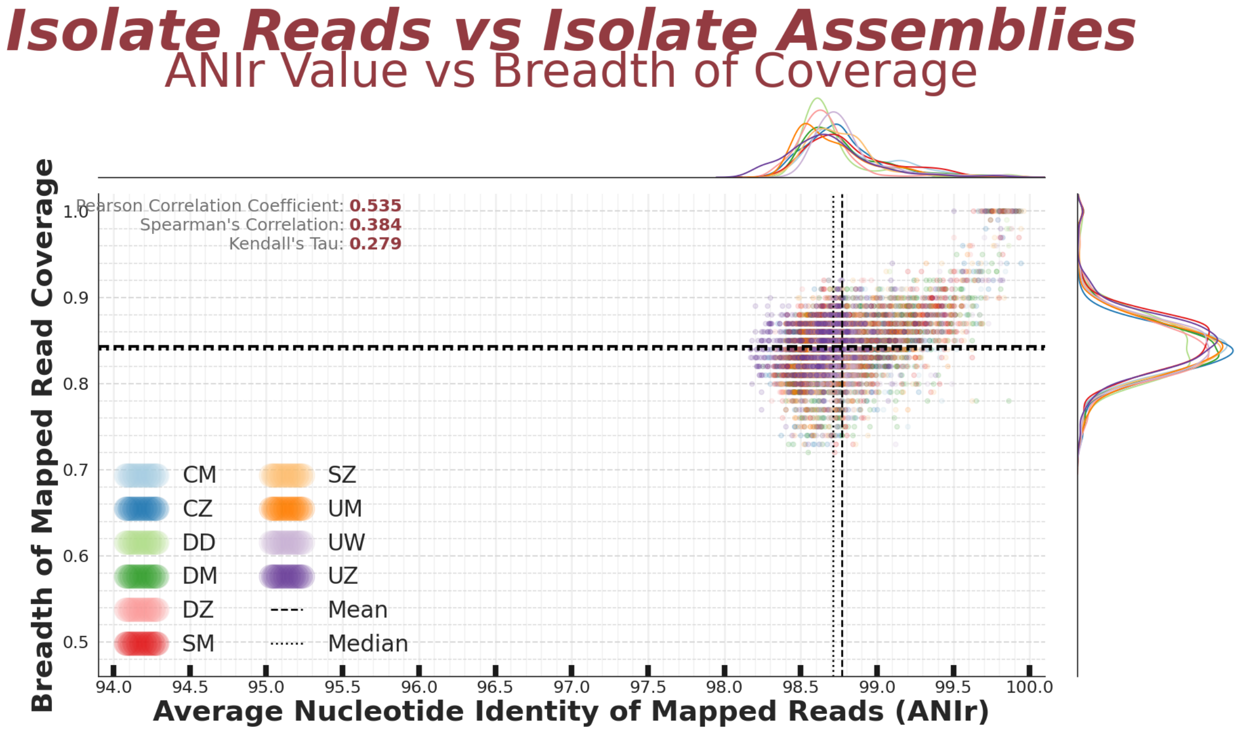


**Supplementary Figure 3: Genomic diversity of *Sal. ruber* isolates assessed by read mapping.** The breadth of coverage for unassembled reads mapped to an assembled genome (y-axis) is plotted against the average nucleotide identity of the mapped reads (ANIr) (x-axis). Datapoints show results of all unassembled reads mapped to all assembled genomes for 102 x 102 draft genomes of the primary *Sal. ruber* population. Colored labels are designations following from Fig. S1 where the first letter is the experimental pond and the second letter is the time point of isolation [Control (C), Dilution (D), and Unshaded-shaded (U) ponds at times Zero (Z), one week (W), and one month (M)]. Kernel density estimates are plotted on the top and to the right of the scatter plot. Note the overlapping distributions and data points show homogeneous genetic diversity between sampling times and ponds. Note that an average of 15.17% of a draft genome is not covered by the reads from another isolate (y-axis breadth), which represent genome-specific genes in the comparison, and the average ANIr value is 98.7%. These results are highly consistent with our pangenome analysis (i.e., genome assembly against genome assembly), showing ~85% of a *Sal. ruber* genome to represent core genes, on average, and the remaining 15% to be accessory genes.

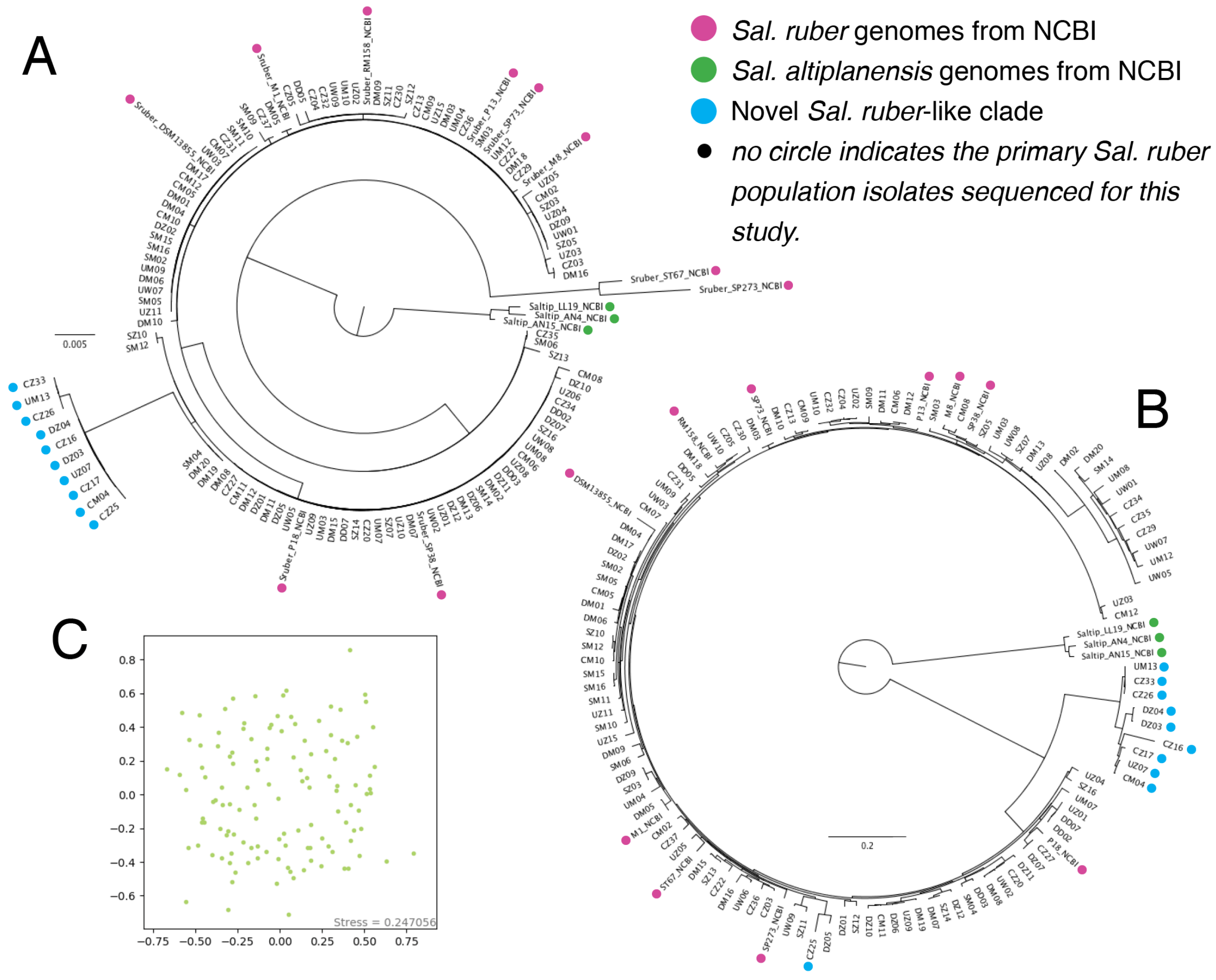


**Supplemental Figure 4: Intra-population genomic diversity**. (**A**) A maximum likelihood phylogenetic tree from RAxML using the full length 16S rRNA gene sequences from 112 *Sal. ruber* isolate draft genomes determined by this study, 10 *Sal. ruber* genomes from NCBI, which were obtained from previous studies in different years and locations (pink), and 3 *Salinibacter altiplanensis* genomes from NCBI (green) as an outgroup. The novel *Sal. ruber*-like clade genomes are denoted in blue. (**B**) An approximately-maximum-likelihood phylogenetic tree from

FastTree using a concatenated set of 106 single copy marker genes of the same genomes as in **A**. (**C**) An NMDS plot from an all vs. all Mash distance matrix computed from draft genomes of the 102 isolates identified as the primary *Sal. ruber* population indicating no clear clustering based on sample time or sample pond.

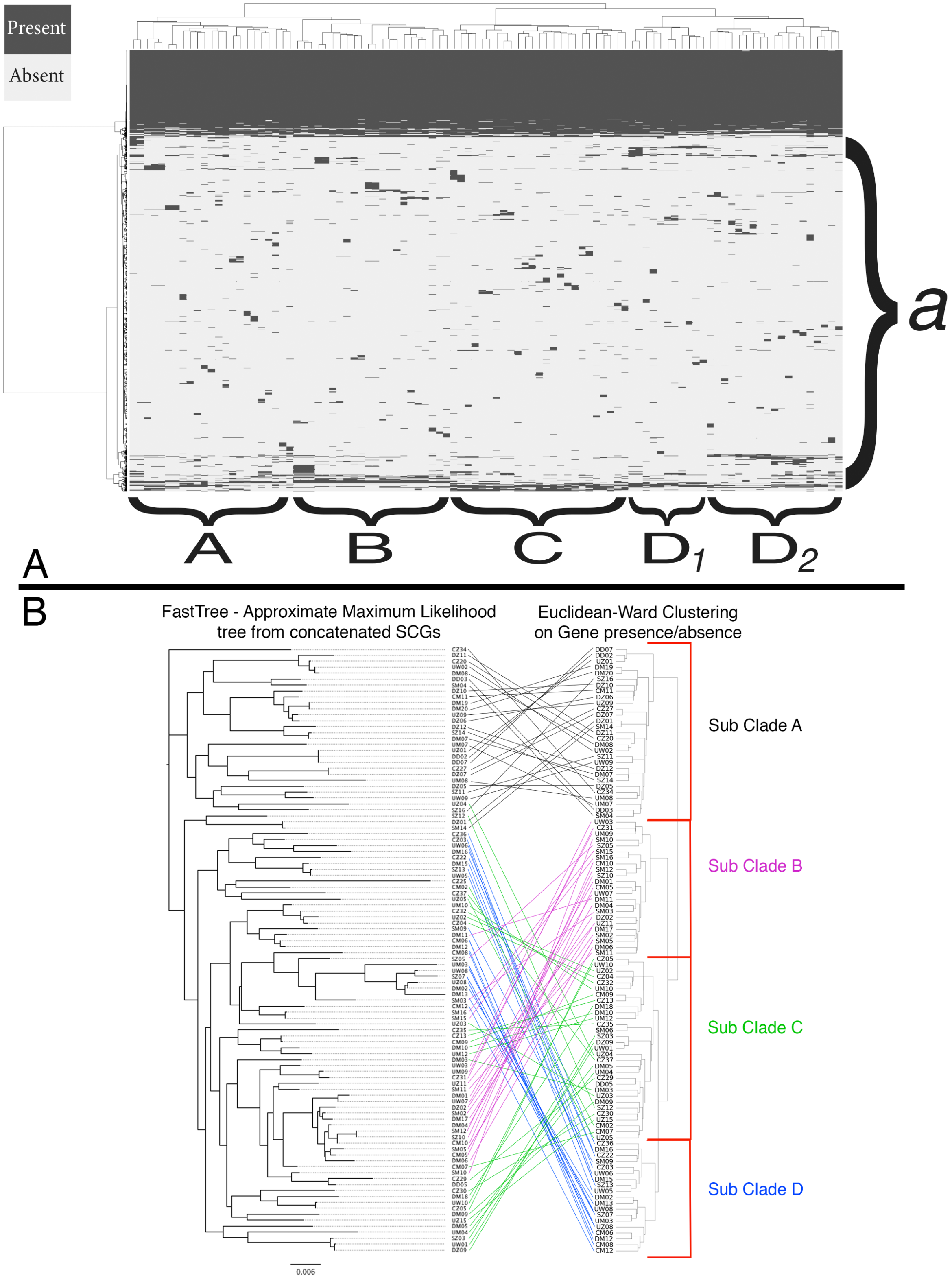
Present
Absent
a
A
B
C
D1
D2
A
B
FastTree - Approximate Maximum Likelihood tree from concatenated SCGs
Euclidean-Ward Clustering on Gene presence/absence
Sub Clade A
Sub Clade B
Sub Clade C
Sub Clade D
0.006

**Supplemental Figure 5: Clustering of *Sal. ruber* isolate genomes based on shared gene content reveals similar but divergent subpopulation structure compared to concatenated gene tree.** (A) Each column represents one *Sal. ruber* draft genome and each row is one non-redundant gene of the *Sal. ruber* pangenome. Genome clustering is based on the pairwise Euclidean distance calculated using the *scipy.spatial.distance.pdistfrom* package and the Ward variance minimization algorithm from the *scipy.cluster.hierarchy.linkage* package in Python. The region designated as *a* highlights the accessory genome and (**A-D**) outline plausible subpopulation clustering (structure). Note that clustering of the genomes based on non-core genes only (as opposed to all genes above) provided essentially the same branching patterns (dendrogram not shown). (B) Comparison of the approximately-maximum-likelihood phylogenetic tree from SCGs with the Ward based dendrogram from Euclidean distance based on the presence or absence of accessory genes. The concatenated alignment of SCGs was stringently trimmed to remove all gap positions and contiguous non-conserved positions of 5 or greater. The lines connect the same gene across both trees.

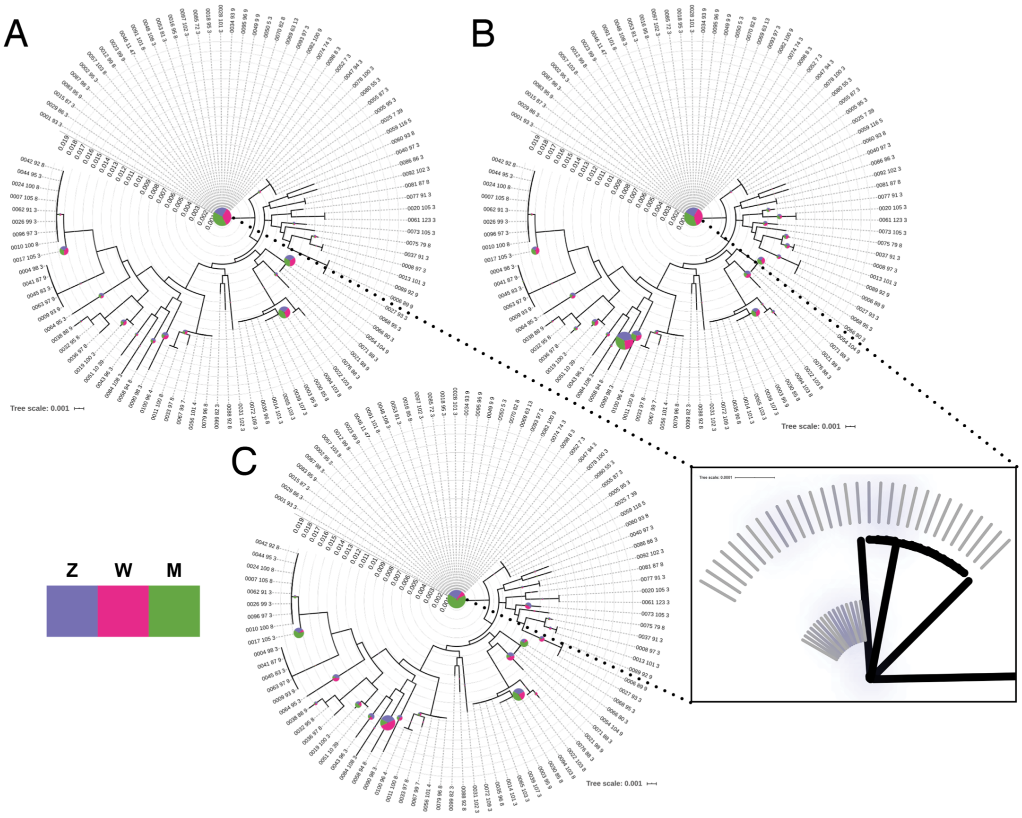


**Supplemental Figure 5: Maximum Likelihood hood trees from rpoB gene sequence alignments with evolutionary read placement.** rpoB sequence for S. ruber draft genomes were aligned with Clustal omega and the tree calculated using RAxML. Metagenomic reads for 3 timepoints each from 3 experimental ponds were aligned and placed with the evolutionary placement algorithm from RAxML. The size of the pie and width of the slice are proportional to the number of reads placed at that position. (**a**) Control pond (**b**) Unshaded-Shaded pond (**c**) Dilution pond

**Supplemental Figure 6: Maximum Likelihood trees of *rpoB* gene sequences of isolate genomes and *rpoB*-carrying metagenomic reads.** *rpoB* sequences from *Sal. ruber* draft genomes were aligned with Clustal omega and the maximum-likelihood phylogenetic tree was built using RAxML. Next, metagenomic reads for three timepoints each from three experimental ponds were aligned to the full-length *rpoB* alignment and then placed on the tree with the evolutionary placement algorithm from RAxML. The size of the pie and width of the slice are proportional to the number of reads placed at that position. (**A**) Control pond (**B**) Unshaded-Shaded pond (**C**) Dilution pond. The color key corresponds to sampling time zero (Z), one week

(W), and one month (M) as in Fig S1. The inset shows the detail of the tree structure obscured by the center pie. Note that several different genotypes (represented by each different pie) were found in each sample, and their relative abundance fluctuated, more or less randomly, across samples (variation in pie and slice sizes) based on the number of short reads uniquely assigned to each genotype.

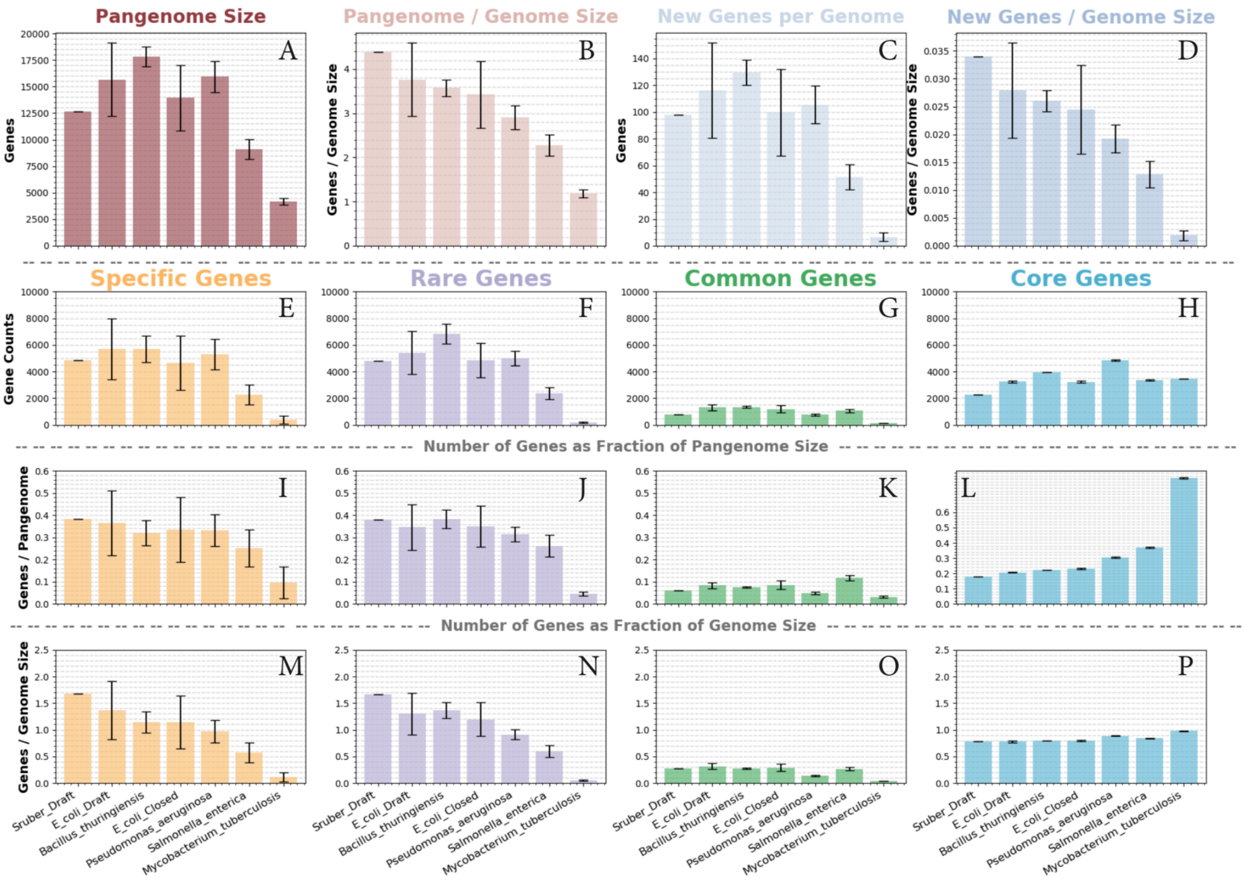


**Supplementary Figure 7: Detailed comparison of pangenome metrics for selected bacterial species.** Pangenome metrics were calculated from draft genome collections for multiple organisms, plus one closed genome collection for *E. coli*. Error bars show the 95% empirical confidence interval of results calculated from 100 random trials each selecting 100 genomes and running 100 permutations. *Sal. ruber* error bars are not shown because only 100 draft genomes from this experiment were available (no random trials). (**A**) Absolute count of total non-redundant genes after the addition of 100 randomly selected genomes. (**B**) Values from (**A**) normalized by the average genome size of each species. (**C**) Mean number of new genes added to the total pan-genome per genome addition. (**D**) Values from (**C**) normalized by the average genome size of each species. (**E-P**) Contribution to the pangenome of different classes of genes (isolate-specific, rare, common, or core) based on their prevalence among the 100 genomes of the species analyzed (see text for details). (**E-H**) The absolute count of genes for each gene class are shown. (**I-L**) Counts from E-H normalized by the total size of the pangenome from (**A**). (**M-P**) Counts from E-H normalized by the average genome size of each of the species analyzed.

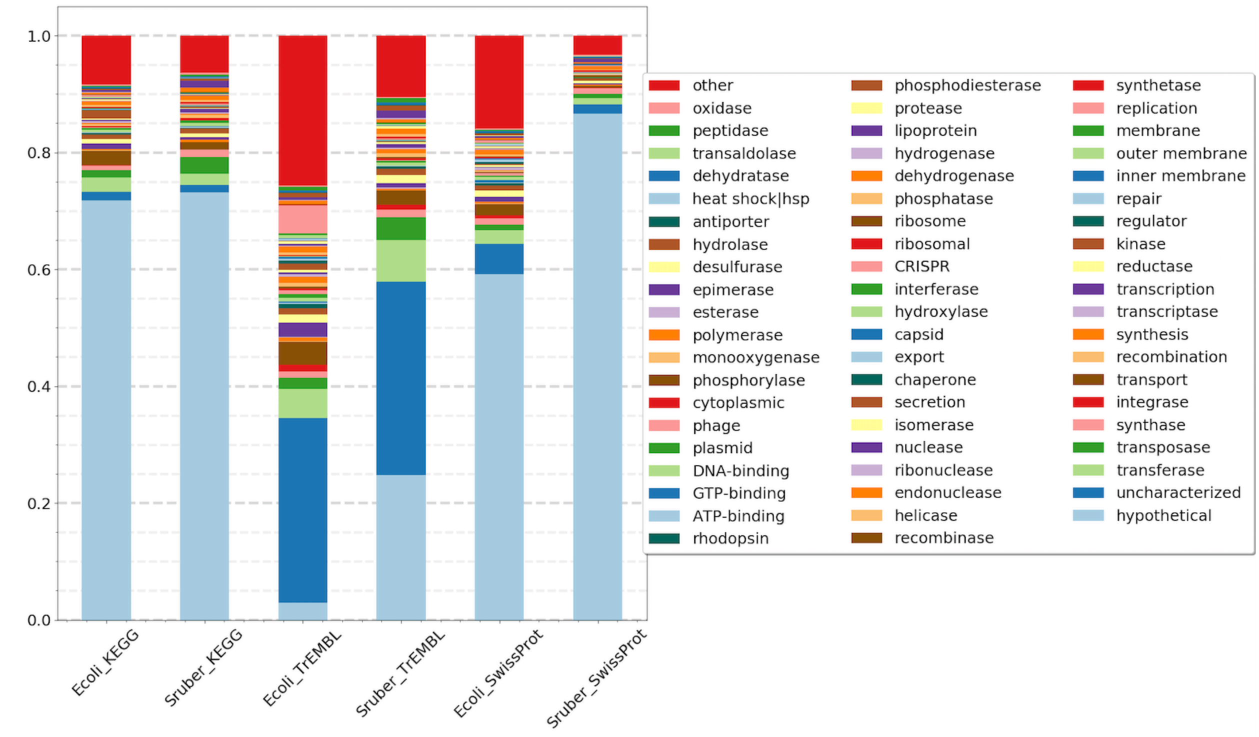


**Supplemental Figure 8: Comparison of the functional gene content of the *E. coli* and *Sal. ruber* pangenomes.** Nonredundant gene clusters from all pangenome experiments were annotated using the KEGG, SwissProt, and TrEMBL databases (see Material and Methods for details). A keyword match was then used against the long gene names of the annotations to generate counts of the functions shown (figure key). Stacked bar plots are normalized to 1 to show the percentage of the total that each category represents.

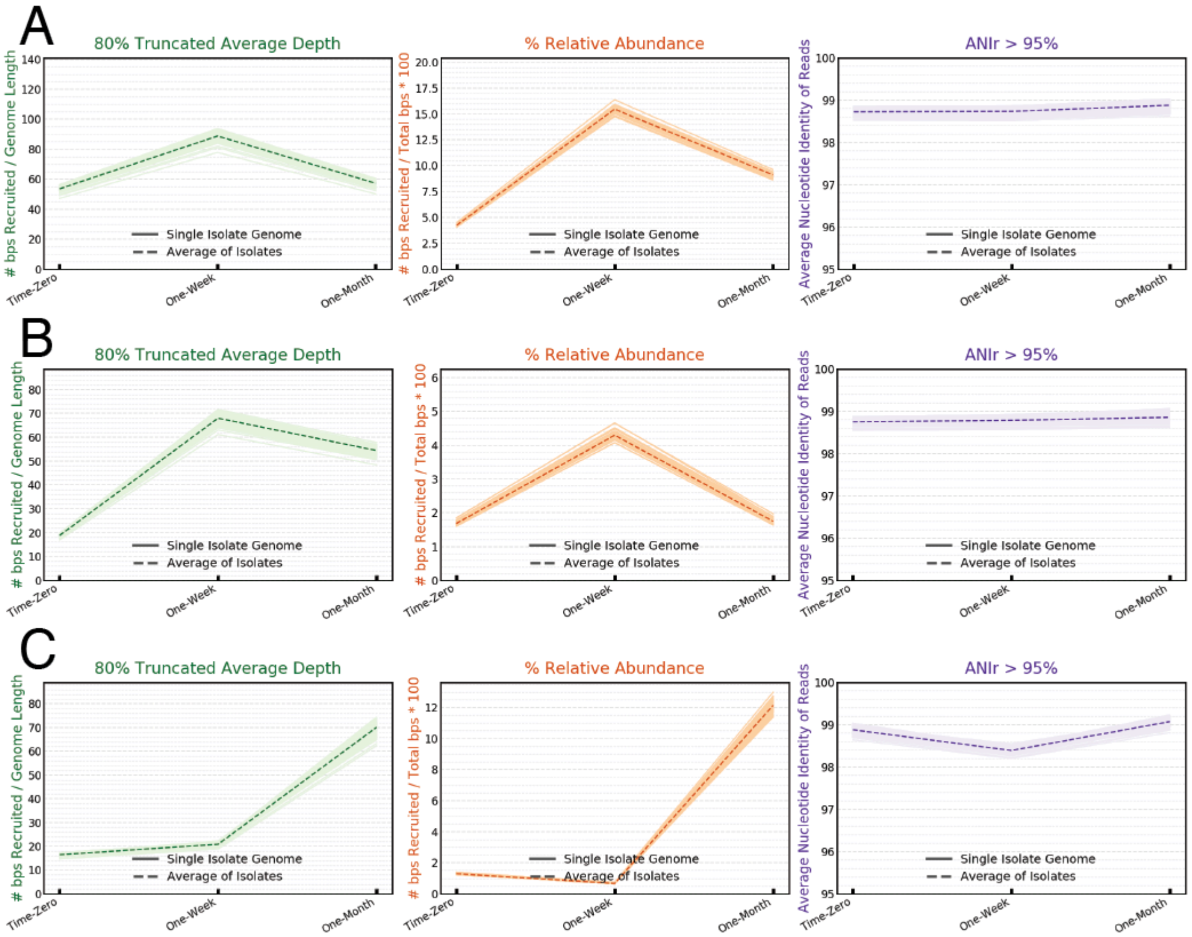


**Supplemental Figure 9: Whole genome abundance dynamics.** Metagenomic reads for three time points each from three experimental ponds were aligned against the *Sal. ruber* isolate draft genomes and TAD80, Relative Abundance and ANIr were calculated and plotted for all *Sal. ruber* isolates individually with the average value indicated. (**A**) Control pond (**B**) Unshaded-Shaded pond (**C**) Diluted pond.

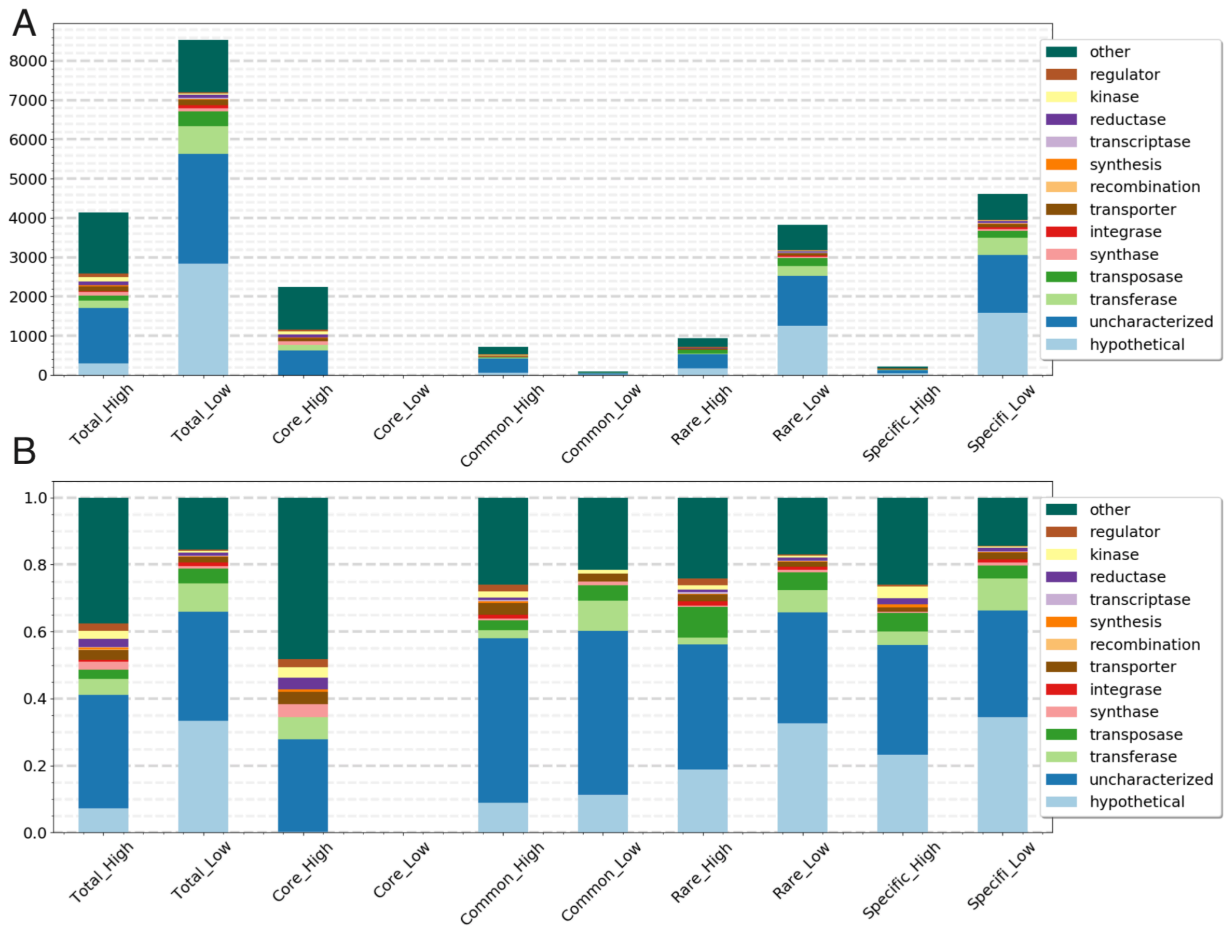


**Supplemental Figure 10: Functional annotation of *Sal. ruber* genes by pangenome gene class and relative *in situ* abundance.** Annotations shown are based on the TrEMBL database. Bar plots are in sets of two with the first (High) counting the genes with an averaged and normalized TAD80 value ≥ 0.25 and the second (Low) counting genes < 0.25. (**a**) Absolute counts and (**b**) Normalized counts.

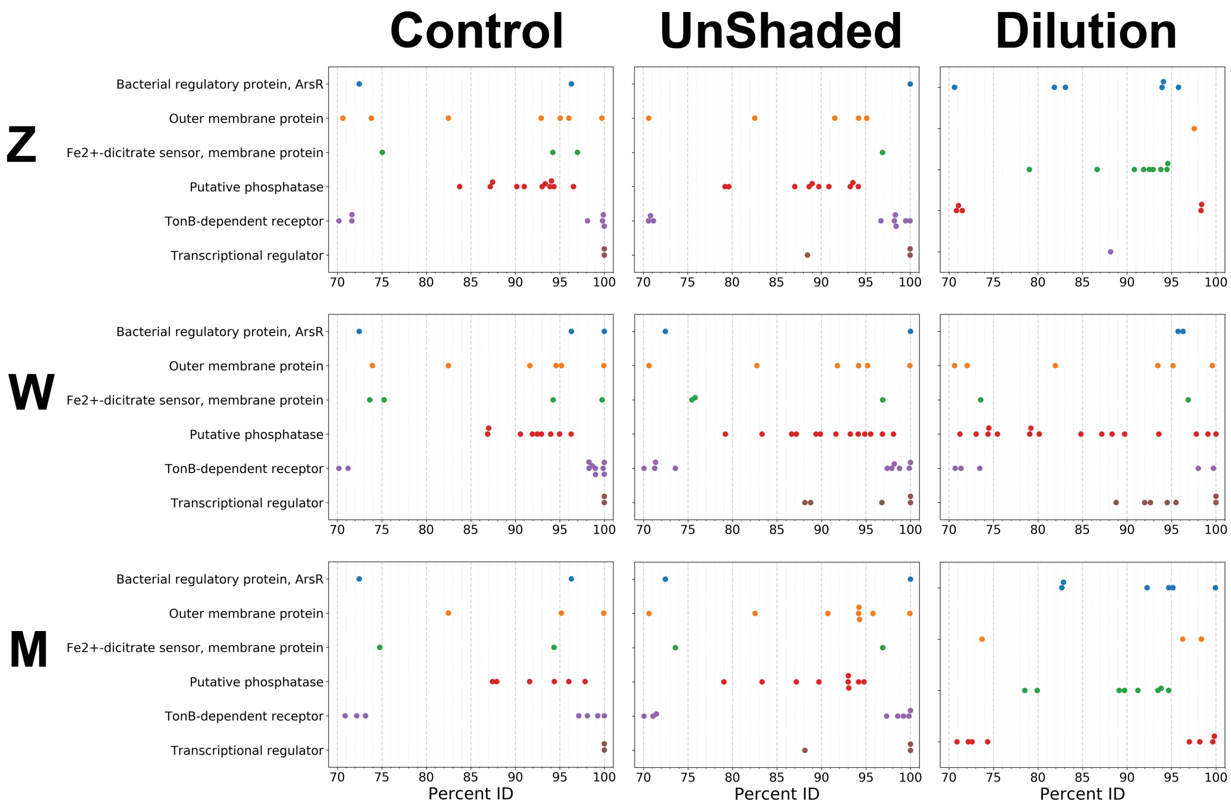


**Supplemental Figure 11: Metagenome assembled contigs matching rare dilution peak genes.** Each dot represents a contig assembled from the metagenome that shared sequence similarity to a rare *Sal. ruber* gene predicted from the draft genome sequence of our isolates. The gene annotation is shown on the y-axis and the percent identity of the gene sequence match to the contig is shown on the x-axis. Genes were selected from the coverage peak identified in the one-weak metagenome sample in the dilution pond series from Figure 4. Columns show the individual ponds and rows shows the time-points. These results show that multiple contigs with high identity matches to *Sal. ruber* were identified in the intermediate salinity metagenomes (23.6% Salts) together with contigs with relatively low identity matches (e.g., <80%), likely indicating that additional community members encode the genes.

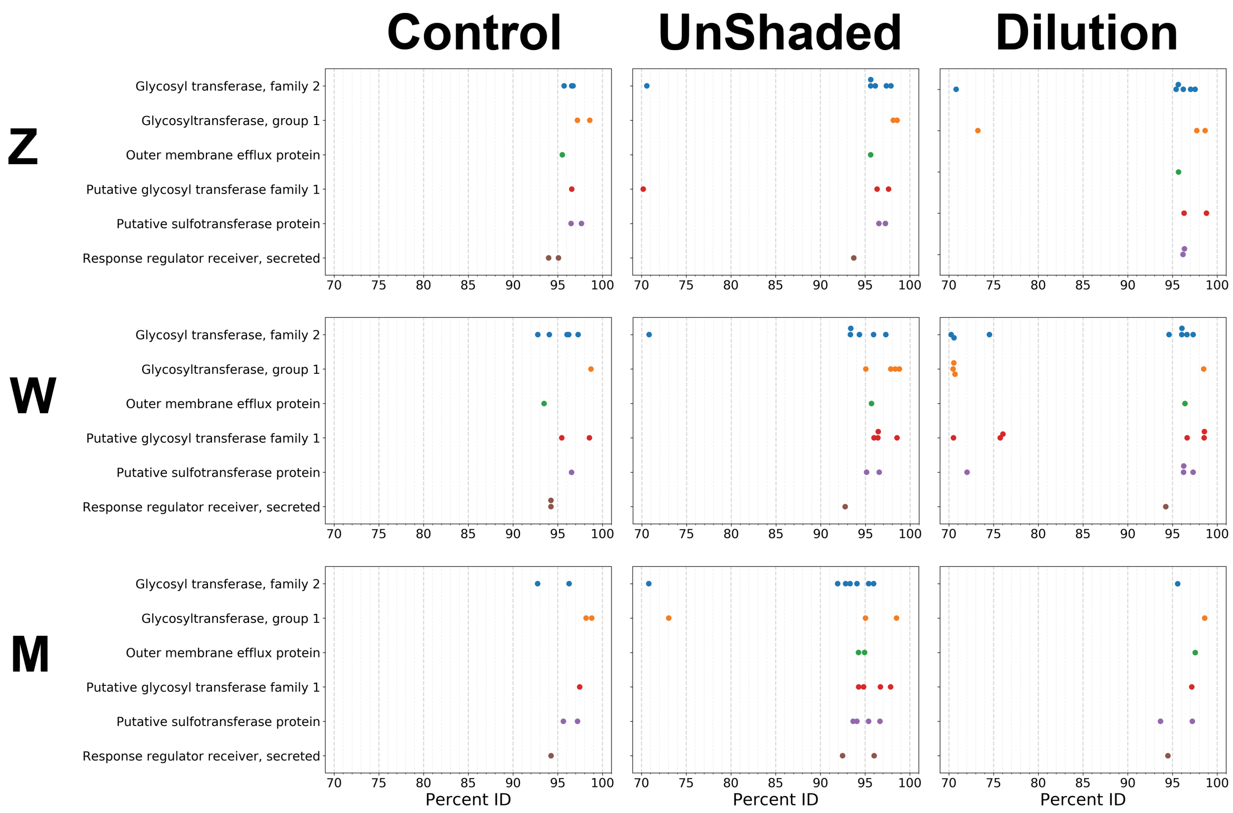


**Supplemental Figure 12: Metagenome assembled contigs matching genome-specific dilution peak genes.** Each dot represents a contig assembled from the metagenome that shared sequence similarity to a genome-specific *Sal. ruber* gene predicted from the draft genome sequence of our isolates. The gene annotation is shown on the y-axis and the percent identity of the gene sequence match to the contig is shown on the x-axis. Genes were selected from the coverage peak identified in the one-week metagenome sample in the dilution pond series from Figure 4. Columns show the individual ponds and rows shows the time-points. These results show that the M8-matching, isolate-specific genes appear to be found only within the *Sal. ruber* species except for in the Dilution T1w where at least one other more distant population other than *Sal. ruber* has similar genes.

| Saltern Pond | Time Point | Length (bp) | Depth (x) | Match (%) |
|---|---|---|---|---|
| Unshaded | T1w | 12 806 | 2.93 | 93.32 |
| Unshaded | T0h | 7 244 | 1.43 | 70.59 |
| Unshaded | T1m | 4 757 | 5.48 | 93.32 |
| Control | T1w | 3 571 | 2.48 | 97.28 |
| Unshaded | T1m | 2 842 | 3.24 | 95.4 |
| Dilution | T1w | 2 823 | 8.17 | 97.31 |
| Unshaded | T0h | 2 385 | 4.67 | 96.14 |
| Dilution | T1w | 2 175 | 17.98 | 70.57 |
| Dilution | T1w | 2 148 | 7.72 | 96.62 |
| Control | T0h | 2 129 | 1.29 | 95.71 |
| Unshaded | T1m | 1 910 | 1.18 | 70.81 |
| Unshaded | T1w | 1 897 | 2.28 | 70.81 |
| Dilution | T1w | 1 574 | 5.10 | 74.53 |
| Unshaded | T1w | 1 454 | 4.93 | 95.91 |
| Dilution | T0h | 1 147 | 2.20 | 70.81 |
| Unshaded | T0h | 979 | 1.97 | 97.38 |
| Dilution | T0h | 975 | 2.10 | 96.21 |
| Unshaded | T0h | 663 | 1.61 | 97.89 |
| Dilution | T0h | 471 | 2.42 | 97.03 |
| Control | T1w | 439 | 1.73 | 94.06 |

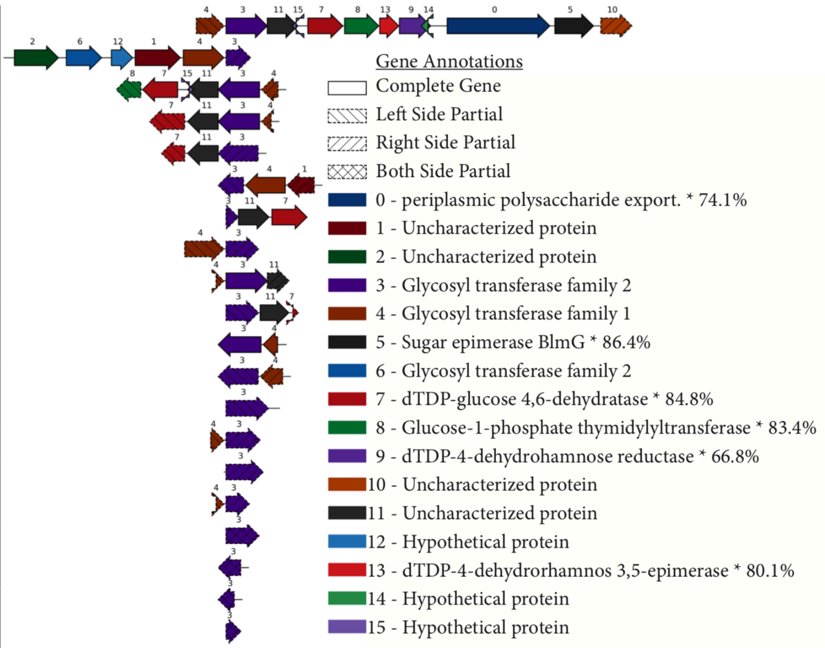


**Supplemental Figure 13: Gene synteny plot of metagenome assembled contigs found to contain isolate-specific genes that peak in abundance in low salinity metagenomes.** Each row represents one contig assembled from one of the 12 metagenomes, and each arrow represents one predicted gene. Contigs were selected if they had a sequence match ≥ 70% nucleotide sequence identity to a glycosyl transferase family 2 gene, which is denoted with the number 3 and color purple. Contigs are centered around the purple gene 3 on the x-axis. The y-axis is sorted by contig length. The table details the corresponding saltern pond, time-point, contig length, sequencing depth supporting the contig, and the percent match of the purple gene 3 to the gene assembled from *Sal. ruber* isolates. An asterisk next to the gene annotation in the legend indicates that *Sal. ruber* was the top sequence match from the NCBI non-redundant database and displays the percent sequence identity of that match. These results show that multiple contigs encoding isolate-specific genes are found in the low salinity metagenome and they are mostly syntenic indicating a shared evolution history.

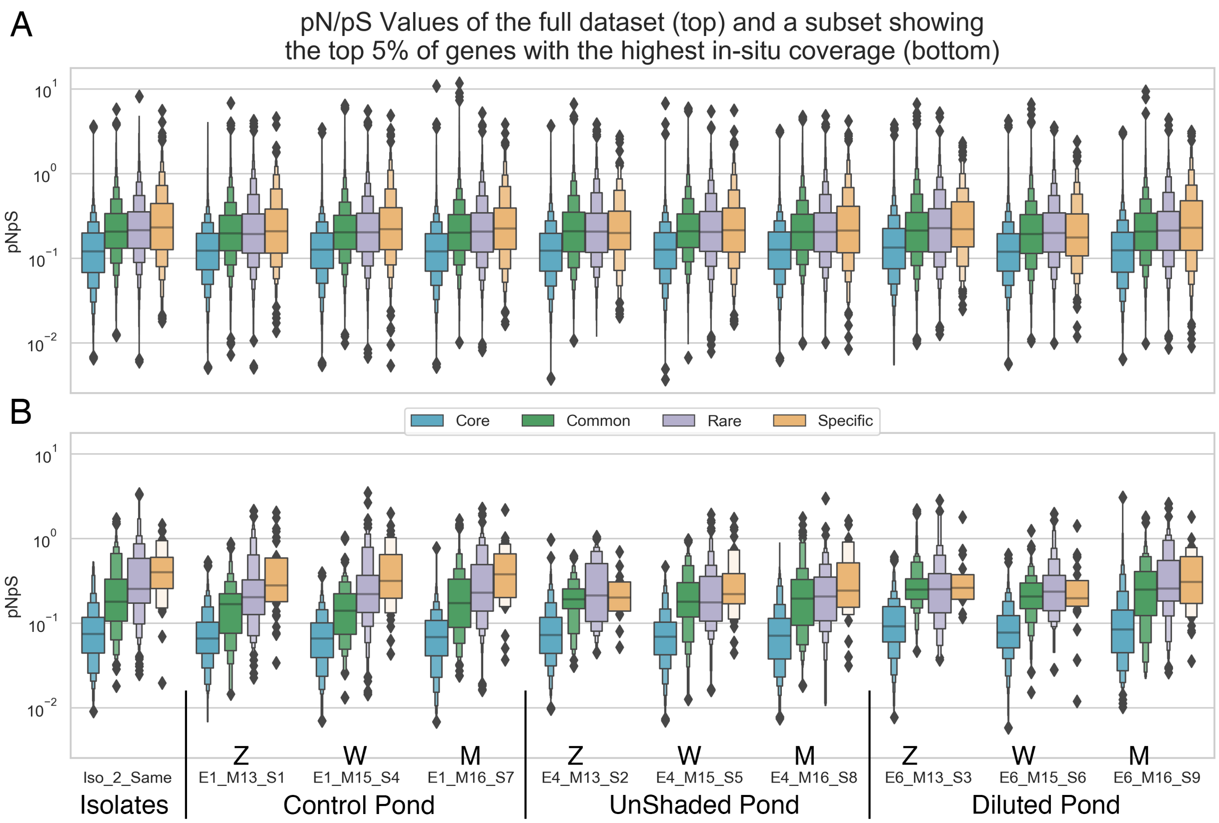


**Supplemental Figure 14: pN/pS values by sample and pangenome gene class.** inStrain (https://instrain.readthedocs.io/en/latest/index.html) was used to calculate pN/pS values from metagenomic reads mapping on isolate genome sequences for the control, unshaded, and diluted pond samples, following program recommendations and using default parameters. To calculate pN/pS from the isolates to compare to the metagenome results, the unassembled reads from all sequenced isolates were combined into a single input and then mapped back to the isolate genome sequences using the same settings as the metagenome mapping. Results were grouped by our defined pangenome gene classes and plotted using letter-value plots (4), showing the distribution with horizontal lines (boxes) representing the median and quantiles for each sample (x-axis) of the pN/pS values for all genes in the *Sal. ruber* pangenome (y-axis) (A) or a subset of the top 5% of genes with highest *in situ* abundance relative to the average whole genome relative abundance (B). Note that the y-axis is in log scale to display the small proportion of genes with pN/pS > 1 ($10^0$), and the great majority of genes show pN/pS < 0.5, regardless of the gene class.


1. Viver T, Orellana LH, Diaz S, Urdiain M, Ramos-Barbero MD, Gonzalez-Pastor JE, et al. Predominance of deterministic microbial community dynamics in salterns exposed to different light intensities. Environ Microbiol. 2019;21(11):4300-15.
2. Viver T, Cifuentes A, Diaz S, Rodriguez-Valdecantos G, Gonzalez B, Anton J, et al. Diversity of extremely halophilic cultivable prokaryotes in Mediterranean, Atlantic and Pacific solar salterns: Evidence that unexplored sites constitute sources of cultivable novelty. Syst Appl Microbiol. 2015;38(4):266-75.
3. Munoz R, Lopez-Lopez A, Urdiain M, Moore ER, Rossello-Mora R. Evaluation of matrix-assisted laser desorption ionization-time of flight whole cell profiles for assessing the cultivable diversity of aerobic and moderately halophilic prokaryotes thriving in solar saltern sediments. Syst Appl Microbiol. 2011;34(1):69-75.
4. Hofmann, H., Wickham, H. & Kafadar, K. Letter-Value Plots: Boxplots for Large Data. Journal of Computational and Graphical Statistics 26, doi:10.1080/10618600.2017.1305277 (2017).
